# LIPTER, a cardiomyocyte-enriched long noncoding RNA, controls cardiac cytoskeletal maturation and is regulated by a cardiomyocyte-specific enhancer

**DOI:** 10.1101/2025.08.10.669544

**Authors:** Svenja Koslowski, Wenhao Zheng, Marouane Benzaki, Michelle Mak, Wang Xiao, Lek Wen Tan, Albert Dashi, Yonglin Zhu, Talal Fawaz, Kenneth Ng, Davy Pham, Francis LeBlanc, Guillaume Lettre, Julie Hussin, Roger Foo, C.J Mick Lee, Anene-Nzelu Chukwuemeka

## Abstract

Cardiac development is characterized by a complex series of molecular, cytoskeletal and electrophysiological changes that guarantee the proper functioning of adult cardiomyocytes (CMs). These changes are defined by cell-type-specific transcriptional rewiring of progenitor cells to form CMs, and are regulated by various epigenetic elements, such as long noncoding RNAs (lncRNAs). LncRNAs are versatile epigenetic regulators as they may act in *cis* or in *trans* to orchestrate important gene programs during cardiac development and may concurrently encode micropeptides. *LIPTER* is one such lncRNA, previously shown to regulate lipid droplet transport in cardiomyocytes and thus an important regulator of cardiomyocyte metabolism. Here we show that *LIPTER* also plays a role in the cytoskeletal maturation of CMs, as loss of *LIPTER* leads to persistent expression of fetal genes, changes in chromatin accessibility, disorganized sarcomeres and impaired calcium homeostasis in CMs. Furthermore, we have identified a cardiomyocyte-specific regulatory enhancer that regulates the expression of *LIPTER* in CMs. CRISPR-mediated inhibition of this enhancer led to reduced *LIPTER* expression in CMs and increased expression of fetal genes. This CM-specific enhancer could therefore be manipulated to control the expression of *LIPTER* for therapeutic benefit. In summary, we have unravelled a novel role of *LIPTER* in CMs’ cytoskeletal maturation and have identified a CM-specific enhancer for *LIPTER*.

## INTRODUCTION

Cardiac development is characterized by a complex series of spatiotemporal and morphological changes culminating in the formation of the heart chambers, cardiac valves and the great vessels^1^. At the cellular level, this involves the differentiation of cardiac progenitor cells (CPC) into the different cell types of the heart, such as cardiomyocytes (CMs), endothelial cells (EC), smooth muscle cells (SMC) and cardiac fibroblasts (CF)^2,3^. Cardiomyocytes, in turn, develop into chamber-specific CMs and undergo electrophysiological, cytoskeletal and metabolic maturation^4^ to become functional adult CMs. These morphological and functional CM changes are defined by extensive transcriptional rewiring, including the upregulation of cell-type-specific myofibrillar and ion-channel genes and the expression of relevant metabolism and calcium handling genes^4–6^. Transcriptional changes are themselves regulated by epigenetic factors such as long noncoding RNA (lncRNA), thus, unravelling novel lncRNA regulators of cardiac development will provide valuable insight into fundamental mechanisms of gene regulation and CM function.

LncRNAs are an important class of epigenetic regulatory elements that orchestrate important gene programs in development and disease^7,8^. They are often cell- and species-specific, consistent with their critical roles in lineage commitment and cell function^7,8^. Several lncRNAs important for cardiac development have been described, such as *Braveheart* (*Bvhrt*), *Heart Brake (HBL1*) and *CARMEN*^9–11^, each acting through unique or diverse context-dependent mechanisms. LncRNAs can act in *cis* to regulate genes nearby or in *trans* to regulate gene expression elsewhere^12^. They may also simultaneously have *cis* and *trans*-regulatory functions and encode functional micro-peptides^12^. Some well-studied mechanisms of action for lncRNAs include recruiting chromatin remodelers to the DNA, interacting with other macromolecules like RNA and proteins or acting as regulatory enhancers^13^. For instance, the lncRNA *Bvhrt* controls the expression of *MesP1* (mesoderm posterior 1), a key regulator of multipotent CPCs, to activate a core cardiovascular gene network^9^. In addition, *Bvhrt* interacts with SUZ12,^9^ a component of Polycomb Repressive Complex 2 (PRC2) and separately interacts with cellular nucleic acid binding protein (CNBP/ZNF9), a zinc-finger protein, to regulate cardiovascular lineage commitment^14^. *HBL1*, another important lncRNA for cardiac development, has distinct roles in the cytoplasm and the nucleus. In the cytoplasm, it interacts with the microRNA miR-1^10^ and the YB-1 protein^15^, while in the nucleus, it recruits the PRC2 complex to various chromatin regions^16^ to regulate different transcriptional programs. For its part, the lncRNA *CARMEN* regulates both CM differentiation^11^ and SMC differentiation^17,18^ through distinct mechanisms. These demonstrate the functional versatility of lncRNAs and underscore the need for deeper studies of the ever-growing list of human lncRNAs^19^.

A CM-enriched lncRNA *LINC00881,* also called *LIPTER,* short for Lipid Droplet Transporter, was recently shown to facilitate the transport of lipid droplets (LDs) to the mitochondria for metabolism^20^. Loss of *LIPTER* did not affect the *in vitro* CM differentiation efficiency of human induced pluripotent stem cells-derived cardiomyocytes (hIPSC-CM)^20^, and there were no changes in the expression of CM-specific genes based on RNA-seq. Nevertheless, upon stressing the hIPSC-CM with either high-glucose media or palmitic acid, there was an accumulation of LDs, mitochondrial damage and apoptosis in CMs lacking *LIPTER*^20^. An earlier study by Liao *et al*, however, used Gapmers to knock down the expression of *LIPTER* in hIPSC-CMs and observed downregulation of important cardiac genes such as *MYH7*, *RYR2* and *CACNA1C,* leading to reduced calcium peaks in the hIPSC-CMs^21^. The authors proposed *LIPTER*’s interaction with SMARCA4, a chromatin remodeler, as the potential mechanism of action^21^. Altered calcium homeostasis with no apparent mitochondrial damage was also observed in another earlier study that used siRNA to knock down *LIPTER* in hIPSC-CMs^22^. This underlines *LIPTER*’s importance in cardiac contractility and points to *LIPTER*’s potentially diverse functions in different contexts. For disease relevance, *LIPTER* is downregulated in dilated cardiomyopathy^23^, ischemic cardiomyopathy^21^ and type 2 diabetes-induced cardiomyopathy^20^. Furthermore, genetic variants associated with gene expression changes of *LIPTER* in the heart have been identified in genome-wide association studies (GWAS) of cardiac rhythm^24^ and right heart structure^25^, amongst others^26^. All these suggest that *LIPTER* plays different important roles in the regulation of cardiac rhythm, structure and metabolism and warrants further study.

To deepen our insight into the roles of *LIPTER* in CM development and function, we analyzed transcriptional and functional changes of *LIPTER*-depleted CMs using two orthogonal models. We performed a CRISPR-mediated deletion of the first exon (referred to as knockout (KO)) and a separate inducible short hairpin-mediated downregulation of *LIPTER* (referred to as knockdown (shRNA-KD)). The inducible shRNA-KD allows us to reduce the expression of *LIPTER* at different time points during CM differentiation to assess its role at different critical stages of CM development. Our data revealed that *LIPTER* plays a role in the cytoskeletal maturation of CMs, as loss of *LIPTER* led to persistent expression of early CM genes such as *MYH6*, *NPPA* and various SMC genes characteristic of immature CMs ^27^. This was accompanied by reduced expression of CM mature genes such as *MYL2*, disorganized sarcomeres and impaired calcium homeostasis. In addition, we identified a CM-specific enhancer that regulates the expression of *LIPTER*. This enhancer contains a variant which is an expression quantitative trait locus (eQTL) for *LIPTER,* and which has been prioritized in a GWAS of cardiac structure and function^25^. This finding is important because the promoter region of *LIPTER* bears H3K27ac and ATAC-seq peaks (features of active promoters and enhancers) in all cardiac cell types, even though the transcript is only expressed in CMs. This suggests that other CM-specific regulatory mechanisms help to ensure its CM-specific expression, and our results point to this CM-specific enhancer as one of such mechanisms. Together, our data provide new evidence of the multiple roles of *LIPTER* in CM development and function and offer novel insight into one of the regulatory mechanisms that regulate its expression in CMs.

## RESULTS

### *LIPTER* is a CM-enriched lncRNA whose expression initiates in early committed cardiomyocytes

Although previous studies have identified *LIPTER* as a CM-enriched lncRNA, here we analyzed its precise temporal pattern of expression using human embryonic stem cell-derived cardiomyocytes (hESC-CM). We employed an established protocol for the differentiation of hESC to CMs through the modulation of the Wnt signalling pathway^28^ (Figure 1A), which yields predominantly ventricular cardiomyocytes. Using this protocol, we performed RNA-seq on eight different time points, which are representative of the stages of CM differentiation *in vitro* and reflective of *in vivo* development (Figure 1A). They are days 0 (hESCs), 1 (meso-endoderm progenitors), 3 (cardiac mesoderm), 5 (multipotent CPCs), 7 (committed CPCs), 10 (early embryonic CMs) and 14 and 28 (fetal CM). For *in vivo* relevance, we analyzed publicly available RNA-seq data performed on human embryonic hearts at different Carnegie stages (CS) of development from CS13 to CS23. At the transcriptional level, CS13 correlates with the day 7 (CPC) stage, while CS16 correlates with the day 14 hESC-CM ^5^. Weighted Correlation Network Analysis (WGCNA) was applied to the RNA-seq data to cluster the differentially expressed genes (DEG) in both datasets into different modules based on their temporal pattern of expression during development (Supplementary Figure 1A). *LIPTER* was found in the turquoise module, which contains mature CM genes like *MYL2* and *MYH7* in both the *in vitro* and *in vivo* datasets (Figure 1B, C). Genes in this module are absent during the pluripotent and mesoderm stages and become expressed as cells transition from the committed CPC (day 7 and CS13) into early CMs (day 10 and CS16), remaining highly expressed afterwards. This module with a mature gene profile clustered differently from other modules, like the yellow module, which contains CM genes characteristic of a fetal gene profile, such as *MYH6 and CARMEN*. The genes in this *MYH6* module are expressed early at the CPC stage and decrease as CMs mature (Figure 1D, E), both *in vivo* and *in vitro*, and are thus indicative of an immature CM gene expression profile. We performed qPCR of *LIPTER*, *MYH6*, *MYL2* and *MYH7* to confirm their temporal pattern of expression (Figure 1F,G), showing that, indeed, *MYH6* expression begins before *LIPTER,* which in turn precedes *MYL2* and *MYH7*. Analysis of the expression of these genes in the *in vivo* dataset also confirmed this trend (Figure 1H). To confirm the CM-specific expression of *LIPTER*, we analyzed cardiac single-cell RNA-seq data from the Human Cell Atlas^29^ and confirmed that it is only expressed in CMs using the UCSC cell browser^30^ (Supplementary Figure 1B). We also performed qPCR analysis of *LIPTER* in our hESC-CMs and commercially available CF, SMCs and ECs to confirm its CM-specific expression (Supplementary Figure 1C). Cellular fractionation of hESC-CM showed a higher abundance of *LIPTER* in the cytoplasmic fraction of the cells (Supplementary Figure 1D), while a 5’ and 3’ Rapid amplification of cDNA ends, identified the transcript with three exons as the most abundant isoform (Supplementary Figure 1E). We also identified another isoform with the same first 2 exons but a truncated third exon (Supplementary Figure 1E).

**Figure 1.**
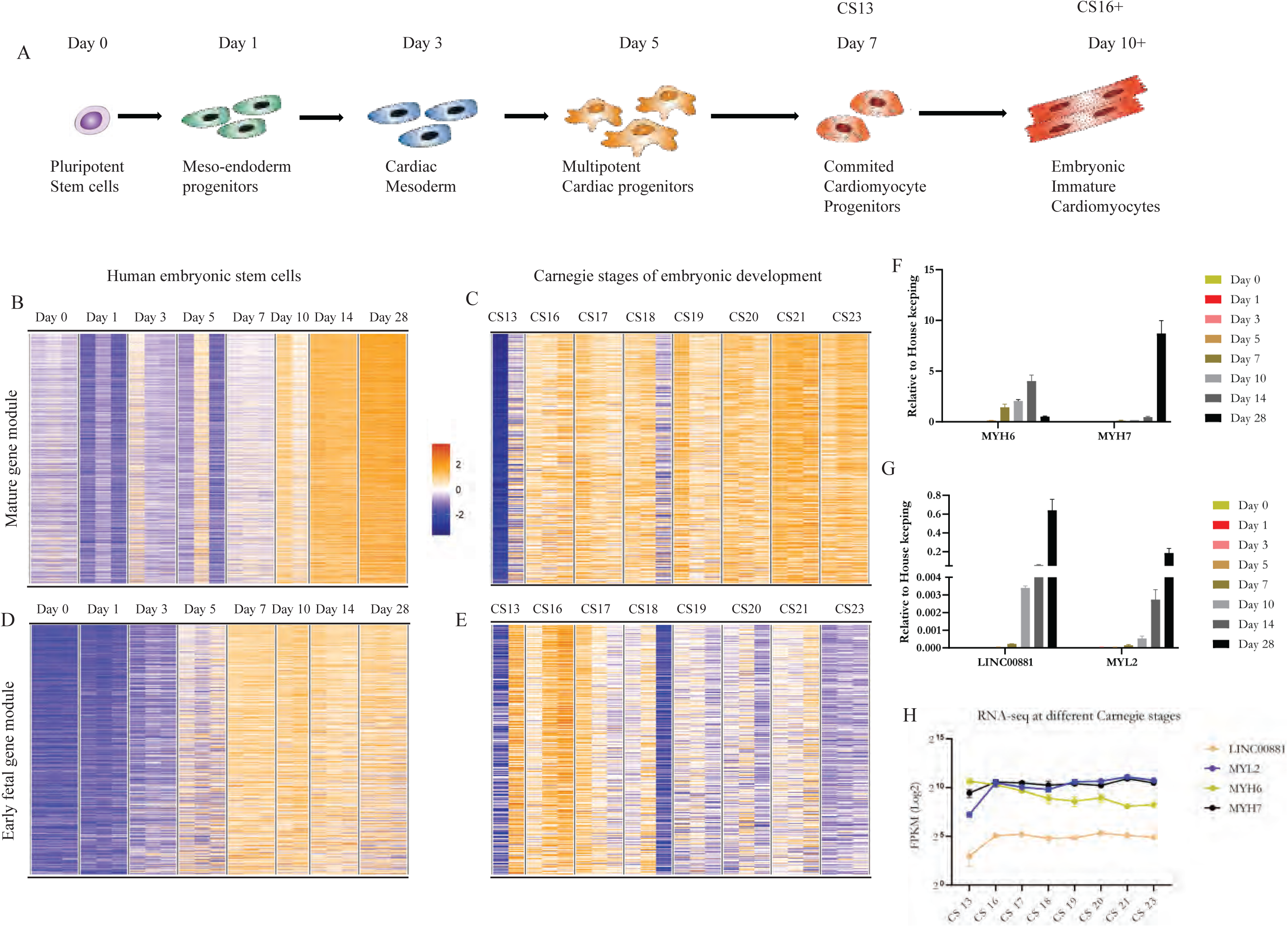
A) A schematic of the differentiation protocol, showing the various developmental stages of the stem cells through the mesoderm stage and cardiac progenitor stage before cardiomyocytes. Prolonged culture leads to increased maturity of the CMs. The corresponding Carnegie stages are listed for Day 7 and Day 10. B, C) A heat map showing the gene expression pattern of genes in the mature gene module, identified by WGCNA, *in vitro* and *in vivo*. D, E) A heat map showing the gene expression pattern of genes in the fetal gene module, identified by WGCNA, *in vitro* and *in vivo*. F, G) QPCR analysis of the temporal expression pattern of *MYH6, MYH7, LIPTER* and *MYL2 in vitro*. H) FPKM values showing the temporal expression pattern of *MYH6, MYH7, LIPTER* and *MYL2 in vivo*.

### Loss of *LIPTER* does not affect CM differentiation efficiency but impedes transcriptional maturation over prolonged culture

To identify *LIPTER*’s roles in CM development, we used both a CRISPR-induced genomic deletion of the first exon and an inducible shRNA KD in a hESC line. This approach with the inducible shRNA allows us to perform knockdown at different timepoints but also helps delineate if the mechanism of action is based on its genomic locus as an enhancer or if the transcript is required for its function. The hESC line has an MYH6-mCerulean3 fluorescent protein (CFP)^31^, which turns cerulean around day 7 of differentiation as the expression of *MYH6* begins. This fluorescence can be observed on the green channel using a fluorescent microscope. Since *MYH6* is an early pan CM gene which decreases in expression as the CMs mature, the CFP intensity also diminishes by the 4^th^ week of culture as the cells mature, with almost no CFP observed by the 6^th^ week (Supplementary Figure 2A). With this line, we generated a monoclonal line with a doxycycline-inducible shRNA to knock down the expression of *LIPTER*. The shRNA inducible construct also has a red fluorescent protein reporter (Supplementary Figure 2B), which turns red upon treatment with doxycycline and indicates the expression of the shRNA (Supplementary Figure 2C). We differentiated the hESCs to CMs^28^ and treated the cells with doxycycline from day 0 of differentiation and every 3 days afterwards to induce and maintain the expression of the shRNA and thus knock down *LIPTER* expression. Upon differentiation, we observed similar levels of CFP in the KD cells during the first 2 weeks (Supplementary Figure 2D), indicating that the loss of *LIPTER* had very little effect on differentiation efficiency, as also evidenced by an earlier study^20^. Given that it was found in the same module as mature genes, we decided to perform long-term culture as a model of *in vitro* maturation to assess its role in CM maturation^4^. The cells were thus left in culture with regular RPMI cell culture media for up to 6 weeks, and we observed that the CFP signal, which indicates the expression of *MYH6,* remained high in the *LIPTER* shRNA-KD CMs (Figure 2A). In contrast, the control cells with the non-targeting shRNA (NTC) that also received doxycycline treatment, and the cells with the shRNA targeting *LIPTER* but without doxycycline treatment, had reduced CFP expression as expected (Figure 2A). We performed a qPCR analysis and observed increased *MYH6* with reduced *MYL2* and *MYH7* in the shRNA-KD CMs compared to the control cells (Figure 2B). Next, we repeated the experiment but started the doxycycline treatment from day 10 (in early CMs after the onset of *LIPTER* expression) and maintained the treatment of doxycycline every 3 days until day 42 (Figure 2C). This was done to assess if there are major differences in the function of *LIPTER* if the KD was performed before the onset of its expression or after. In addition, we generated two knockout (KO) clones by deleting the promoter and first exon of *LIPTER* (Supplementary Figure 2E) using the same MYH6-CFP hESC line and differentiated these cells into CMs.

**Figure 2.**
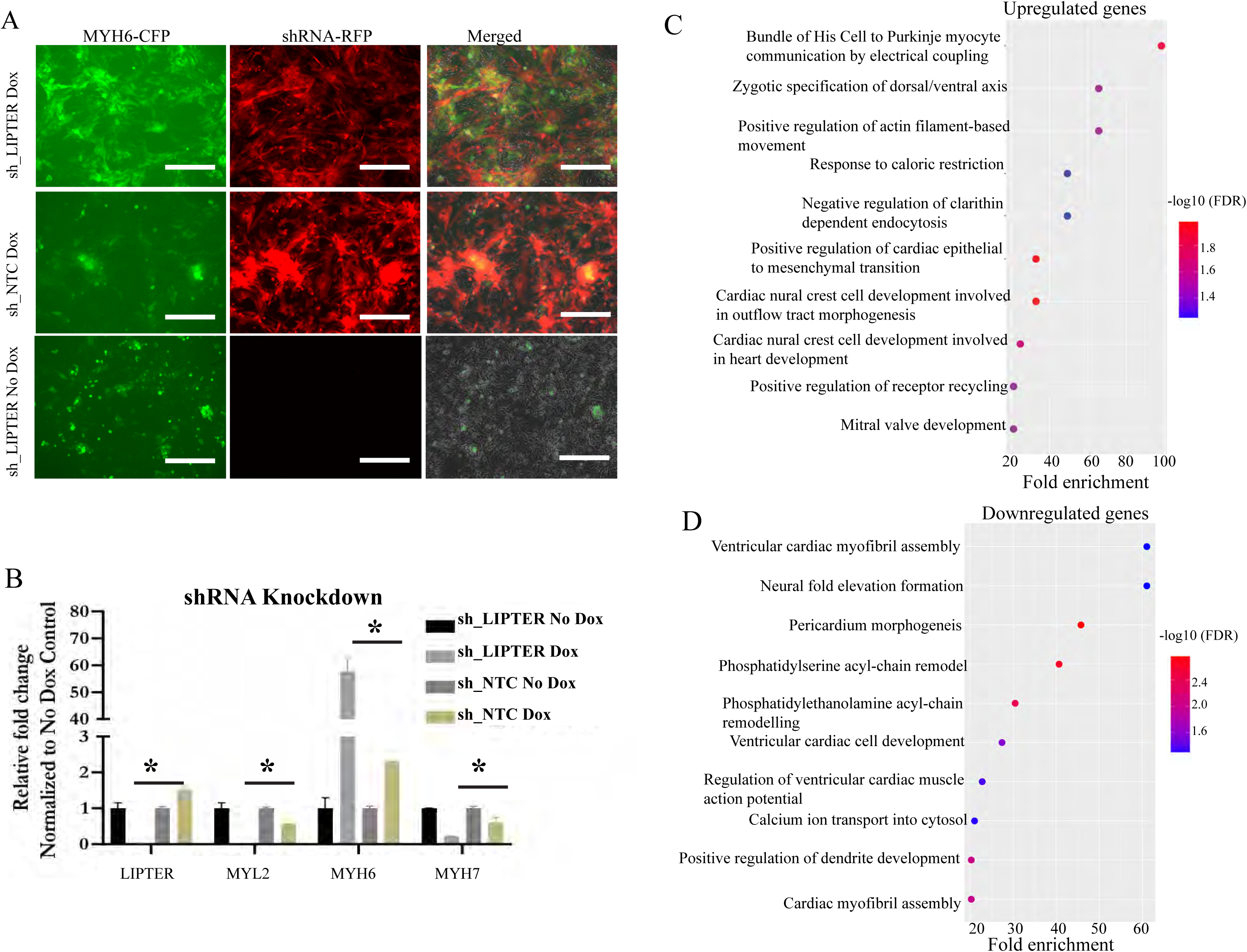
A) Fluorescent images showing CFP intensity indicative of MYH6 and RFP intensity indicative of the shRNA. There was increased CFP in the CMs with the knockdown of *LIPTER*. B) A qPCR validation showing increased *MYH6* and reduced *MYL2* and *MYH7* in the CMs with knockdown of *LIPTER*. 3 biological replicates were used for each sample, * P < 0.05. C) Gene ontology analysis of shared upregulated genes. D) Gene ontology analysis of shared downregulated genes.

We observed similar results of higher CFP intensity in the KO cells, confirmed with qPCR (Supplementary Figure 2F), and thus, we performed RNA-seq on day 42 of these CMs for a total of three main groups, namely: 1) Control and shRNA KD with doxycycline from day 0 (two biological replicates each), 2) Control and shRNA with doxycycline from day 10 (three biological replicates each), 3) Two separate knockout clones of the *LIPTER* with a shared WT control group (three biological replicates each) (Supplementary Figure 3 A-D). From the RNA-seq, first, we analyzed each group individually to identify differentially expressed genes and gene ontology (GO) terms and then compared the results to identify common GO terms. For the downregulated genes, using the AmiGO GO^32^ tool, common terms were “Heart Development,” “Cardiac contraction,” “Myofibril Assembly,” and “Cytoskeletal Organization,”. For the upregulated genes, the common themes included “Extracellular matrix organization,” “Heart development,” “Artery and aortic valve morphogenesis,” and “Lipid metabolism,” amongst others (Supplementary Tables 1-8). Next, we compared all datasets to highlight the shared upregulated and downregulated genes for a total of 288 downregulated and 239 upregulated genes in both shRNA-KDs and the knockout samples (Supplementary Table 9). Gene ontology analysis for the shared upregulated genes, included terms such as “Bundle of His cell to purkinje myocyte communication”, “outflow tract morphogenesis” (Figure 2C, Supplementary Table 10). Some genes in this group include *MYH6, NPPA, ACTA2*, and *FBN1*. These are early embryonic CMs and smooth-muscle cell (SMC) genes, which are also expressed in early CMs. For the downregulated genes, AmiGO showed terms like Cardiac myofibril assembly, phosphatidylserine acyl-chain remodelling, and Actin Filament Organization (Figure 2D, Supplementary Table 11). Some genes in this group included *LIPTER, MYL2, MYL3, OBSCN, SORBS1*, and *LMOD3,* amongst others. These are genes involved in myofibrillar and cytoskeletal organization ^4,33^.

Next, we compared our RNA-seq result with the previously published study on *LIPTER* by Han *et al*^20^. From their data, there were 15 downregulated genes and 117 upregulated genes after the deletion of *LIPTER* (supplementary Table 2 from their study) ^20^. We only had 2 overlaps, *LIPTER* and *COBL,* for the downregulated genes, while there were 7 overlaps from the upregulated genes, including *MYLK, FBN1, ITGA1*, and *MMP2*. We also downloaded and analyzed the RNA-seq from the earlier study by Liao *et al*^21^, in which the authors performed a 3-day Gapmer KD study, and obtained 792 upregulated and 637 downregulated genes. We compared our shared RNA-seq data with their RNA-seq data using FDR <0.05 and observed 37 and 29 overlaps for downregulated genes and upregulated genes, respectively. Some overlapping downregulated genes of interest between the Gapmer study and our study include *RYR2, CACNA1C, SORBS1* and *BMP7*, genes involved in cardiac contraction and development. Comparing Liao *et al*’s study to the study by Han *et al*, there were 2 overlapping downregulated genes (*LIPTER* and *GLP1R*) and 5 overlapping upregulated genes. Given the little overlap, we reasoned that while the exact genes may not be the same, the ensemble of genes may be involved in the same process. Thus, using AmiGO, we performed GO analysis for the 117 upregulated genes from Han *et al’s* paper and 792 from Liao *et al*’s study, and compared them to our study to identify common terms. Common GO terms were “Aorta development”, “Angiogenesis” and “Aorta smooth muscle cells” (Supplementary Table 12-13). We also used the Enrichr tool^34^ to identify the most likely cell type associated with the upregulated genes from our study, Han *et al* and Liao *et al*, and the top cell type for all datasets was Smooth Muscle Cells based on transcriptomic data from the Human Gene Atlas. Indeed, despite the few precise gene overlaps, the gene profile for upregulated genes had various SMC cytoskeletal genes and extracellular matrix genes in both studies. This includes matricellular proteins *CCN1, CCN2* and gamma actin gene *ACTG2* in the study by Han *et al* ^20^. Similarly, our study had upregulation of genes like *CNN3, MYH11* and *ACTA2*. Given the very few downregulated genes from the Han *et al* study, there was no significant cell type associated with the gene set from their study. Together, this suggests that loss of *LIPTER* may lead to changes in the cytoskeletal organization, impeding CM maturity, and favouring the expression of SMC genes.

### Loss of *LIPTER* leads to extensive chromatin accessibility changes in cardiomyocytes

Given the transcriptomic changes observed in CMs upon the loss of *LIPTER*, we reasoned that the analysis of the chromatin changes in *LIPTER*-depleted CMs would help identify the transcription factors that may be driving these changes and thus provide insight into *LIPTER*’s mechanism of action. Thus, we performed the assay for transposase-accessible chromatin (ATAC-seq) on shRNA KD cells and controls. Although the shRNA system enables assessment of chromatin dynamics across different time points, for this study, we generated data with *LIPTER* KD from day 0. First, we performed a principal component analysis (PCA) on the six samples (3 controls and 3 KD) and observed that the controls and KD clustered separately (Supplementary Figure 4A). A heatmap analysis also showed a clear separation between the control and the KD cells (Supplementary Figure 4B). Distribution of the ATAC peaks with respect to distance from the transcriptional start site (TSS) was similar in all groups (Supplementary Figure 4C). Next, we performed a differential analysis of the accessible chromatin regions and identified over 600 regions that were differentially accessible in each direction (Figure 3A). These included *MYH6* locus in the upregulated regions (Figure 3B). Most of the changes occurred in distal regions located > 50kb from the nearest TSS (Figure 3 C,D).

**Figure 3.**
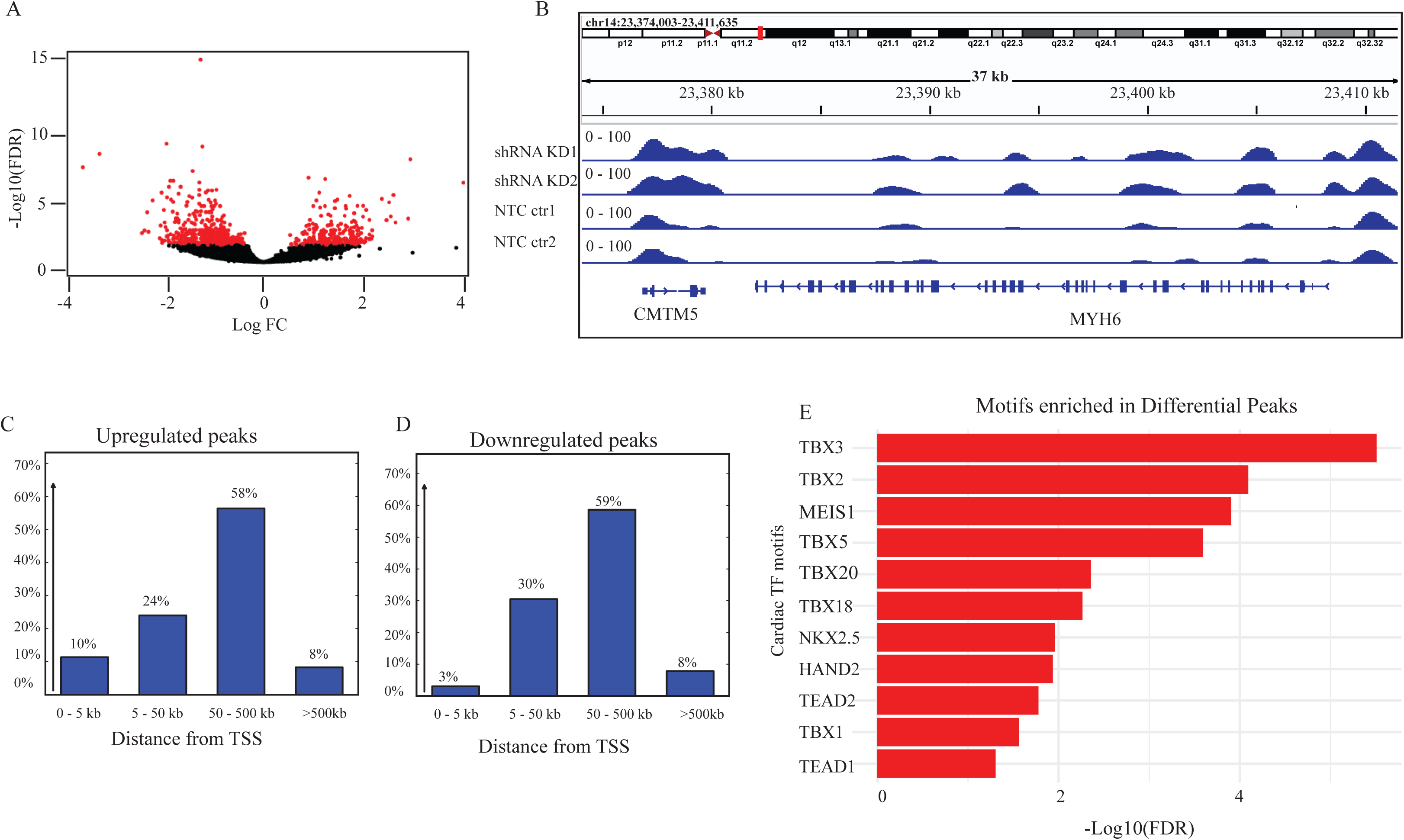
A) Volcano plot showing the differentially accessible chromatin regions based on DESeq2 with FDR < 0.05. B) An IGV plot of the *MYH6* locus, with higher ATAC peaks in the shRNA KD CMs. C,D) Distribution of the differentially accessible chromatin regions for the upregulated and downregulated peaks, respectively. E) Enriched transcription factor motifs in the differentially accessible chromatin regions using JASPAR2020.

Genome Regions Enrichment of Annotations Tool (GREAT) GO analysis, which associates genomic regions with genes^35^, identified “Visceral Muscle development”, “Cardiac atrium morphogenesis” and “Cardiac cell development” as top-enriched terms of genes linked to regions with increased accessibility (Supplementary Table 14). This is consistent with a more immature cardiac gene profile observed in RNA-seq, as many atrial genes and visceral or smooth muscle cell genes are expressed early in ventricular CMs before being downregulated. For downregulated regions, the top gene ontology terms included “Heparan sulphate proteoglycan catabolic process” and “regulation of skeletal muscle differentiation” (Table 15). Heparan sulphates are important regulators of multiple signalling pathways and have been implicated in fibroblast growth factor (FGF) and TGF beta signalling in the heart^36^. Indeed, Perlecan, a heparin sulphate proteoglycan, was recently shown to be important for cardiac development^37^. Perlecan-depleted CMs differentiated efficiently *in vitro*; however, long-term culture (over 30 days) led to structural, contractile, metabolic, and ECM gene dysregulation in these CMs^37^.

We performed motif analysis of the differential peaks from our ATAC-seq and identified the TBX family of transcription factors (TBX3, TBX2, TBX5, TBX20 and TBX18), MEIS1, NKX2.5 and HAND2 as being enriched in these peaks (Figure 3E). Of particular interest is the TBX2-TBX20 axis, given their role in CM specification and the downregulation of TBX20 in our RNA-seq data. TBX20 plays multiple roles in cardiac development ^38,39^, including promoting chamber CMs and suppressing and confining TBX2 to cells of the atrioventricular canal and outflow tract^40^. TBX2, on the other hand, suppresses the formation of chamber CMs and instead directs myocardial cells towards cells of the atrioventricular canal and outflow tract ^40–42^. TBX20 also cooperates with members of the TGF beta signalling pathways like the bone morphogenetic proteins (BMP) and FGF family, to exert its role in cardiomyocyte differentiation ^43,44^. These results point towards a potential mechanism of action through cooperation with TBX20.

### Loss of *LIPTER* leads to disorganized sarcomeres, impaired calcium homeostasis and electrophysiological changes in the CMs

Our gene ontology results from the RNA-seq indicated the downregulation of genes involved in myofibril assembly; thus, we proceeded to analyze sarcomere organization in the KO and KD cells. We performed immunostaining of ACTC1 (cardiac actinin) and observed more defined sarcomere organization and more consistent cell morphology in the wild-type cells compared to the KO. In contrast, the KO cells had a more stress-fibre-like arrangement of the sarcomeres (Figure 4A-C). We used the Sotatool^45^ to analyze the sarcomere organization, and indeed, the analysis revealed a higher sarcomere organization score in the control cells compared to the shRNA KD and CRISPR-mediated KO cells (Figure 4D,E). Next, we performed a patch-clamp to analyze the electrophysiological properties of the CMs. For this analysis, we only used the CRISPR-mediated knockout CMs to avoid possible interference by doxycycline, given the known positive and negative effects of doxycycline on cardiac electrophysiology ^46,47^. Our results showed a tendency to reduced action potential amplitude, action potential duration and a reduced upstroke velocity in the KO cells (Figure 5 A-E). Although the reduced action potential amplitude and duration weren’t significant, these features are characteristic of immature cardiomyocytes. The analysis of calcium current density, however, showed that the ratio of current (pA) over cell capacitance (pF) was lower in WT than in the KO (Figure 5F,G), indicating a higher calcium channel density in the control CMs^48,49^. This also points to a more mature calcium homeostasis in the control cells and confirms previous studies on impaired calcium homeostasis upon downregulation of *LIPTER*.

**Figure 4.**
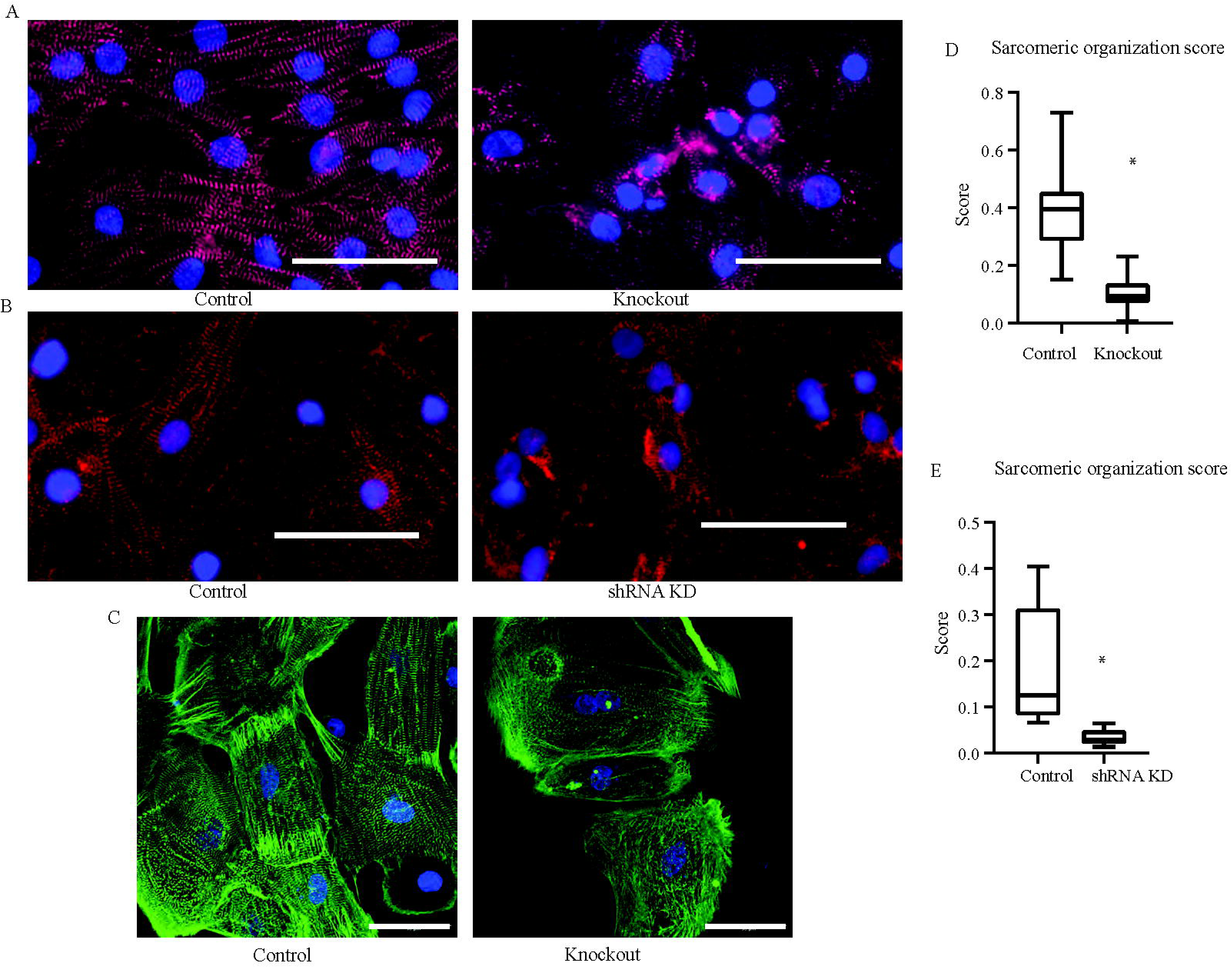
A) Cardiac actinin immunostaining in control and knockout cells showing increased sarcomeric disorganization in the cells with depletion of *LIPTER*. B) Cardiac actinin immunostaining in control and shRNA cells showing increased sarcomeric disorganization in the cells with depletion of *LIPTER*. C) Cardiac actinin immunostaining in control and knockout cells showing the increased stress-fibre-like organization in cells with depletion of *LIPTER.* D,E) Quantification of the sarcomeric organization using the Sota tool. Scale bar 50 μm. * = P value < 0.05

**Figure 5.**
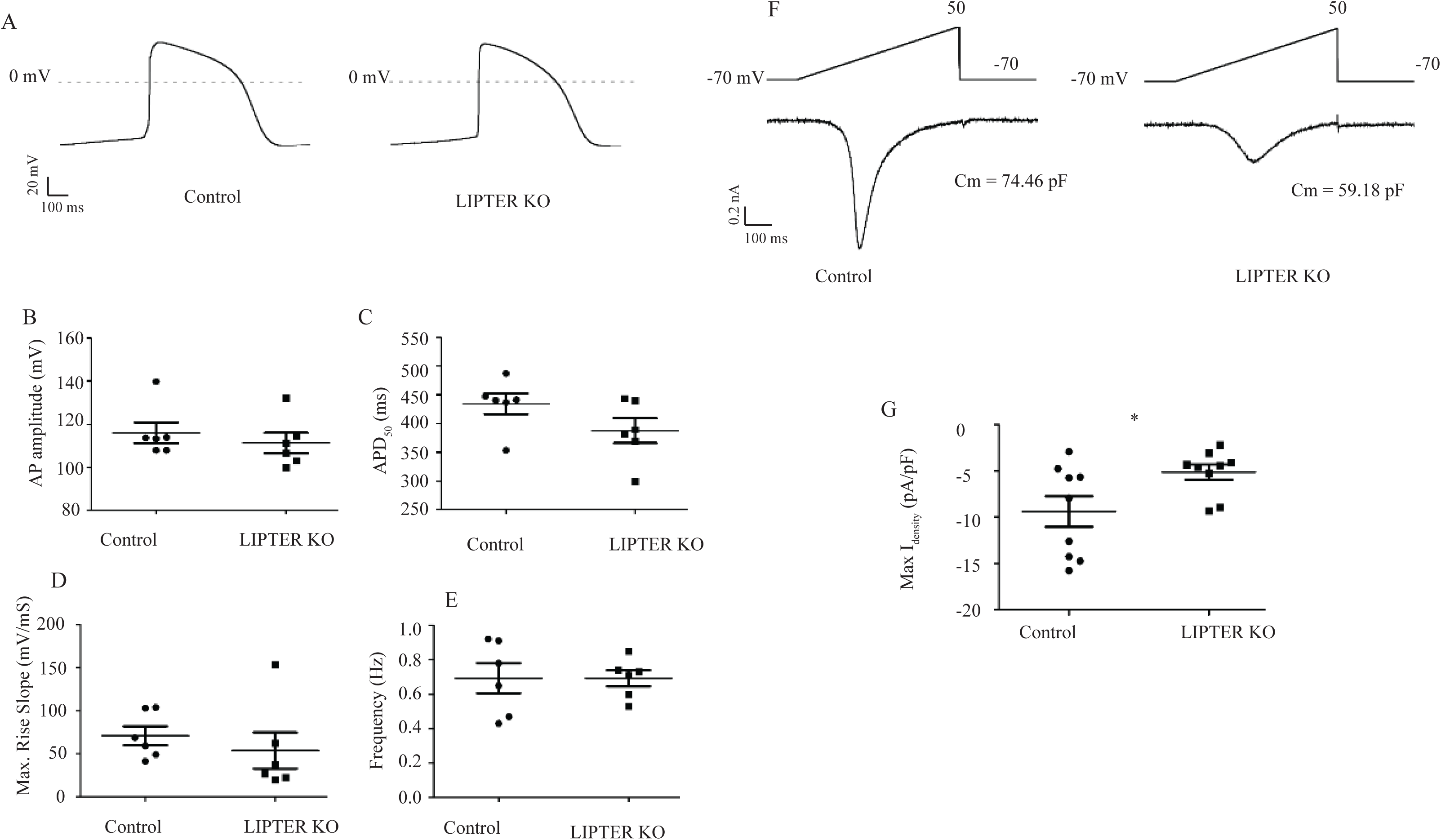
A) A representative trace of the action potential in the control and knockout CMs. Quantification of the B) action potential amplitude, C) action potential duration, D) Maximum rise slope and E) Frequency in the control and knockout CMs. F) Representative trace of the calcium current density. G) Quantification of the maximum calcium current density in control and knockout CMs. * = P value < 0.05

### A CM-specific enhancer contributes to LIPTER expression in CM

An interesting observation from our analysis of publicly available single-cell ATAC^50^ (www.catlas.org) and RNA-seq was that the promoter region of *LIPTER* has ATAC-seq peaks in all cell types of the heart (Supplementary Figure 5A), even though the transcript is only expressed in CMs (Supplementary Figure 1A). We confirmed this using an already published in-house snATAC and snRNA-seq^51^ of the heart (Figure 6A). We also analyzed our in-house bulk H3K27ac ChIP-seq and ATAC-seq of hESC-CM and commercially available cardiac fibroblasts, endothelial cells and smooth muscle cells and indeed observed ATAC-seq and H3K27ac ChIP-seq peaks in all these cell types (Figure 6B). Furthermore, publicly available H3K27ac and H3K4me1 ChIP-seq (https://chip-atlas.org/peak_browser) reveal that the promoter region has H3K27ac and H3K4me1 marks in all cells in the heart. ReMAP^52^, a high-quality catalogue of ChIP-seq of hundreds of transcription factors (TF) in humans, shows the binding of various TFs at the *LIPTER* promoter in different cardiac cell types. This includes CTCF, P300, SS18-SRX in cardiac fibroblasts, CTCF, ERG, FOS in endothelial cells, TBX5, MED1, and GATA4 in CMs. These findings suggest that although *LIPTER* expression is CM-specific, the promoter region may act as a regulatory enhancer for other genes in non-CMs.

**Figure 6.**
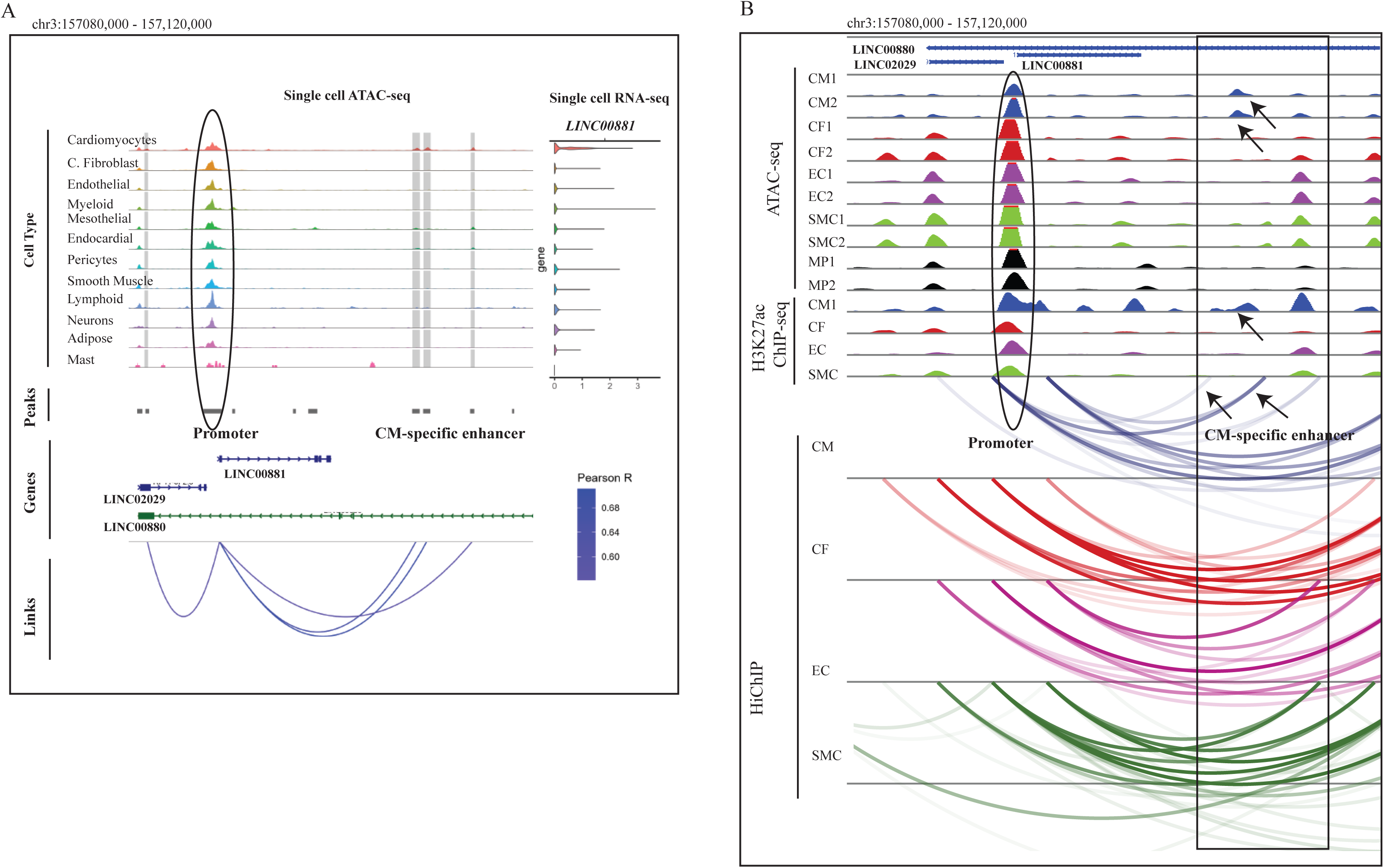
A) Single cell ATAC-seq (top left) and single cell RNA-seq (right) peaks of *LIPTER*’s genomic region from previously published data^51^ to identify the various cell types of the heart. The black oval highlights the promoter region of *LIPTER* and shows that it is accessible in all cell types of the heart. The grey boxes show the cardiomyocyte-specific enhancer linked to *LIPTER*. B) A panel showing our bulk ATAC-seq, ChIP-seq and HiChIP of different cells of the heart. The black oval highlights the promoter region of *LIPTER* and shows both ATAC and H3K27ac ChIP-seq peaks in all cells of the heart. The black box and arrows indicate the CM-specific enhancer and the CM-specific HiChIP links.

We thus reasoned that there may be additional CM-specific regulatory mechanisms that help restrict *LIPTER* expression to CMs; hence, we interrogated our previously published activity-by-contact-derived (ABC) enhancer map of the human heart ^53^. The ABC is a model to predict enhancer-promoter (EP) regulation and assumes that an enhancer’s effect on a gene is a combination of the enhancer activity (ATAC-seq, H3K27ac ChIP-seq) and how often it contacts the gene (Hi-C)^54^. Our ABC analysis on the human heart identified the top two associated regulatory enhancers, (Supplementary Figure 5A, Figure 6B black box), but of the two, one was CM-specific with a chromosomal location at chr3: 157,108,500 – 157,111,135. The other enhancer was also present in endothelial cells, even though the peak was higher in CMs. The CM-specific enhancer also had CM-specific HiChIP interactions not found in other cell types (Figure 6B, arrows). Furthermore, employing the correlation analysis using the Pearson R method to link regulatory regions to genes^55^, this same CM-specific enhancer was identified as one of the enhancers linked to *LIPTER* expression in CMs, confirming our ABC analysis (Figure 6A, blue links). MED1, which binds to this region as highlighted in the ReMAP catalogue^52^ is known to bind to enhancers and facilitates EP interactions^56^. In addition, NKX2.5, ESRRG, TBX5 and GATA4, which also bind to this region in CMs based on the ReMAP catalogue, are TFs important for CM development and function ^22^, confirming that the region is indeed an enhancer in CMs. The only other cell type that appears to have multiple TF binding on this region from the ReMAP catalogue is K562 cells, which are transformed cancer cell lines. Furthermore, a variant rs11928162 located in this region is an eQTL for *LIPTER* in the heart and has been identified as a significant GWAS SNP in studies about genetic determinants of right heart structure and electrophysiological traits ^25,57,58^.

To show that the enhancer regulates *LIPTER* expression, we performed a CRISPR inhibition (CRISPRi) experiment using a hESC line that expresses the KRAB-dCas9. KRAB recruits a heterochromatin-forming complex that leads to histone methylation, deacetylation and gene repression. First, we designed two guide RNAs (gRNA) targeting the enhancer region, cloned them into a lentiviral vector, transduced the CRISPRi hESCs, differentiated the cells into CMs and performed qPCR. Our qPCR result showed a 40 −50% reduction in the expression of the transcript using both gRNAs, however only one was statistically significant. (Supplementary Figure 5B). For long-term culture, we chose the one that was statistically significant and performed qPCR on day 42 and observed a similar gene expression pattern as the KO cells, reduced *LIPTER* and *MYL2,* with increased *MYH6* and *NPPA* (Figure 7A). Next, to analyze the cardiac specificity of the enhancer activity *in vivo,* we PCR amplified the enhancer sequence and cloned it upstream of a minimal promoter and GFP reporter (minP-eGFP) plasmid. We produced AAV9 particles containing the construct, and as a control, we used the heart-specific chicken cardiac troponin T (cTnT) promoter coupled to the same minP-gGFP construct. The AAV particles were injected into the thoracic cavity of p10 mice, and three weeks post-injection, the heart and liver tissues were harvested since AAV9 has cardiac and liver tropism^59^. RNA was extracted from the tissues, and qPCR was performed using primers for GFP. Our results show that the full enhancer sequence (Full sequence), like the chicken cTNT promoter, led to a 4-fold higher GFP expression in the heart compared to the liver (Figure 7C). Finally, to assess which fragments of the enhancer are the key drivers of the enhancer activity, we fragmented the enhancer sequence into four parts broadly based on the ENCODE cCRE annotations. We cloned these individual fragments upstream of the minP-eGFP as described above and performed the same experiment highlighted above. While two of the fragments had a higher GFP expression in the heart than in the liver, the abundance of GFP was lower than with the full enhancer sequence, suggesting a cooperativity of the TFs within the enhancer (Figure 7C).

**Figure 7.**
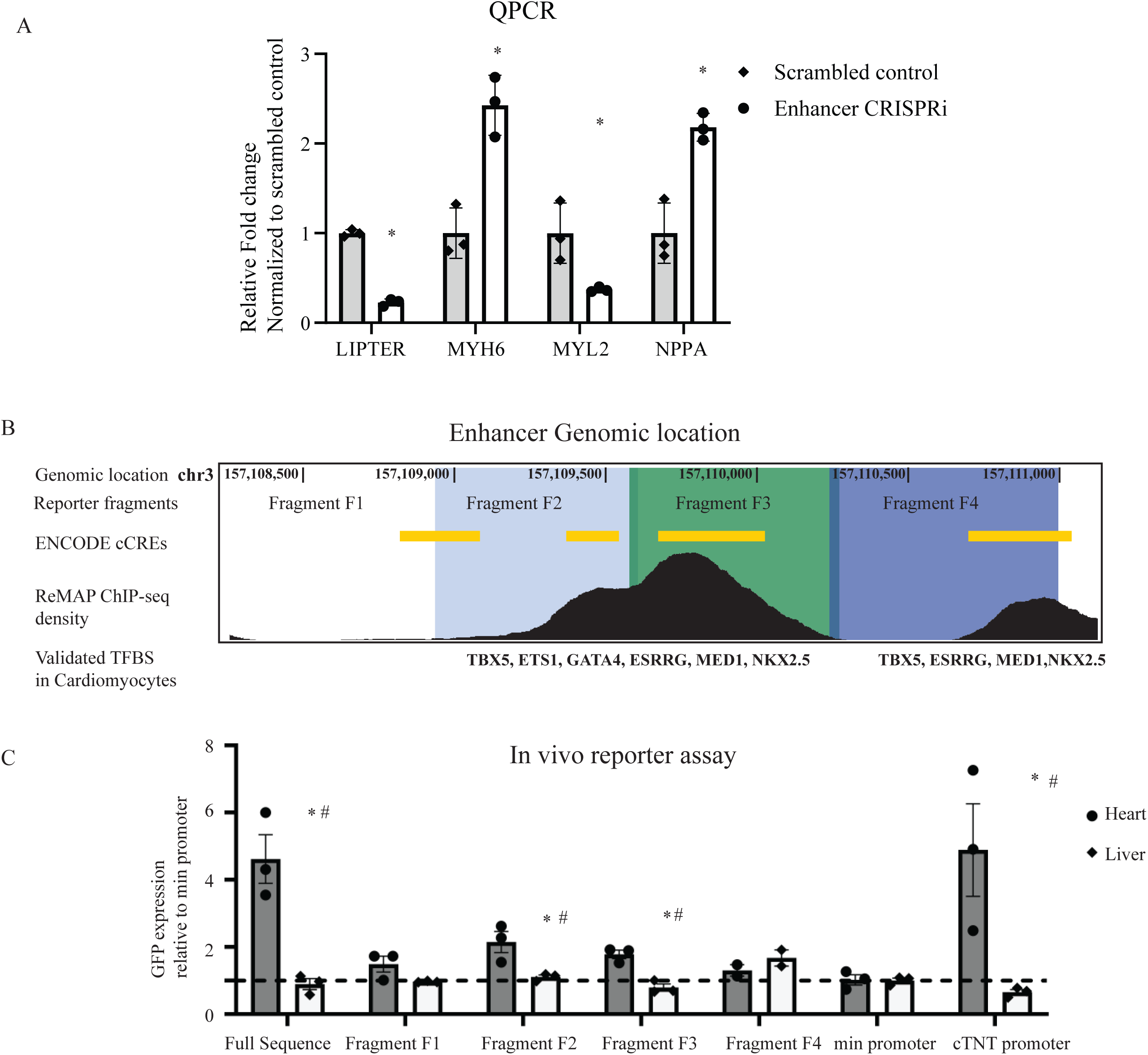
A) QPCR analysis of the expression of *MYH6, NPPA, LIPTER* and *MYL2* after enhancer CRISPRi in the hESC-CMs compared to the scrambled control. There is increased *MYH6* and *NPPA* with reduced *MYL2* and *LIPTER*. * = P value < 0.05. B) UCSC genome browser screenshot of the genomic location of the CM-specific enhancer, identifying the ENCODE-identified cis-regulatory regions and the ReMAP-identified transcription factor binding sites. It also highlights the fragments used for the *in vivo* reporter assay. C) QPCR analysis of GFP expression after the *in vivo* reporter assay in mice. There is a higher abundance of GFP transcript in the mouse hearts compared to the liver. * = P value < 0.05 compared to minimal promoter, # = P value < 0.05 compared to the liver.

## DISCUSSION

Cardiovascular diseases are a major cause of morbidity and mortality from the cradle^60^ to the grave^61^, thus, unravelling the regulators of cardiac development and function is a critical first step in identifying potential therapeutic targets. LncRNAs are versatile regulators of different biological processes, and while some have specific functions, several lncRNAs play different roles and have different mechanisms of action in different contexts. Here we show that loss of *LIPTER* leads to compromised electrophysiological and cytoskeletal maturation of cardiomyocytes in long-term cell culture. These forms of long-term cell culture of CMs over 35 days lead to ultrastructural and electrophysiological maturation of hESC-CMs^62,63^, thus, this model allows for the analysis of the role of *LIPTER* in cardiac maturation. There is a direct relationship between cytoskeletal organization and LD transport, as LDs are transported on cytoskeletal fibres by motor molecules like myosins^64,65^. Destabilizing the cytoskeleton can lead to LD accumulation^65,66^, while alterations to the LD metabolism can lead to cytoskeletal disorganization^64^. Furthermore, one of the gene ontology terms in the downregulated genes was phosphatidylserine acyl-chain remodelling. These acyl chains are fatty acid tails attached to the glycerol backbone of phosphatidylserines. Phosphatidylserines are major components of cell membranes and can play a role in lipid organization and trafficking, suggesting that there is an underlying disturbance in lipid metabolism, which may become evident upon lipid stress to the CMs. This raises exciting new questions in unravelling the role of *LIPTER* in CM development and function. Is the LD accumulation a consequence of cytoskeletal disorganization or a cause? Are both processes distinct but related phenomena co-regulated by *LIPTER*? Interestingly, the protein pulldown experiment by Han *et al* identified mostly cytoskeletal and ribosomal proteins^20^ associated with *LIPTER*. Although the authors only validated MYH10, given its known association with lipid droplets, their mass spectrometry data identified ACTC1, MYH9, and MYL6 as proteins associated with *LIPTER*. Furthermore, the only other overlap for the downregulated genes of their RNA-seq with our RNA-seq is *COBL*, an actin nucleator known to play an important role in the organization of the actin cytoskeleton^67^. This supports our data that *LIPTER* plays a role in the cytoskeletal organization and calls for further analysis of its other mechanisms of action.

Besides the cytoskeletal changes, we also observed impaired calcium homeostasis, altered electrophysiological properties and extensive transcriptional and chromatin accessibility changes. The changes in calcium homeostasis have been observed in earlier studies^21,22^. In addition, genetic studies have linked variants in *LIPTER* to changes in cardiac electrical conduction and cardiac structure ^24,25,57,58,68^. This provides further evidence of the role of *LIPTER* in cardiac contractile function and structure, which may be independent but linked to its role in the transportation of LDs. The increase in SMC genes and genes involved in extracellular matrix development is also of great interest in understanding the role of *LIPTER* in cardiac development. CMs and SMCs have similar developmental origins and diverge during development^2^, and the existence of a bipotent CM and SMC cardiac progenitor cell has been described by various studies^18,27^. Furthermore, smooth muscle cell actin (*ACTA2*) is expressed early in CM and is downregulated as CMs mature^69^. Another evidence of their shared developmental pathways is that a myocardial-to-vascular smooth muscle cell transition is one of the processes involved in the development of the cardiac outflow tract^70^. This transition is characterized by the reduction of expression of CM markers like *MYL2* and the gain of SMC markers like *ACTA2* by CMs. Increased *HES1* expression and reduced *TBX20* expression are hallmarks of this transition process, and these transcription factors have been proposed as the regulators of this transition. Indeed, one of the GO terms of upregulated genes in our RNA-seq is “outflow tract morphogenesis”, and interestingly, we find similar TF changes in our RNA-seq. There was increased *HES1* in all four RNA-seq datasets and reduced *TBX20* in both KO lines and shRNA-treated cells from day 0, but there was no significant change in the cells where KD of *LIPTER* began from day 10^70^. As highlighted above, TBX20 plays a role in maintaining the identity of chamber cardiomyocytes (ventricular and atrial cardiomyocytes) while suppressing the identity of outflow tract myocardial cells, which express many SMC genes. These results suggest a developmental and specification role of *LIPTER* beyond its role in LD metabolism and warrant further studies to delineate its full spectrum of action.

While our RNA-seq and ATAC-seq point to a potential TBX20 pathway through which *LIPTER* exerts its role, another potential mechanism of action not explored in this study is through a micropeptide encoded by *LIPTER*. Indeed, *LIPTER* possesses a small open reading frame (smORF) that encodes a predicted micropeptide of 5kDa. However, analysis of its coding potential has produced conflicting results so far. Riboseq data from human hearts identified *LIPTER* as a transcript associated with ribosomes, while an *in vitro* translation experiment observed that a micropeptide of about 5 kDa is translated from the *LIPTER* transcript^71^. Similarly, a mass spectrometry analysis of the proteome from the human heart identified a peptide fragment that matches the predicted peptide from the *LIPTER* transcript^72^. However, Han *et al* ^20^ did not observe any peptide in the western blot experiment performed in their overexpression experiment in HEK cells^20^, and a mass spectrometry analysis from the study that performed the *in vitro* translation did not identify any such peptide either ^71^. A more detailed study, such as overexpressing the *LIPTER* transcript but with a mutated start codon (ATG) in a knockout cell, may help to tease apart its various roles and mechanisms of action and identify if the smORF is required for its function.

Finally, we have identified a CM-specific regulatory enhancer which contributes to the CM-specific expression of *LIPTER*. This chromatin region is only accessible in CMs and harbors variants which are expression quantitative trait loci for *LIPTER*. Manipulation of this CM-specific enhancer can therefore be used to modulate the expression of *LIPTER* for therapeutic purposes. Indeed, upregulation of *LIPTER* has been shown to rescue cardiac function in mice with high-fat diet-induced cardiomyopathy and type 2 diabetes-induced cardiomyopathy, making it a promising therapeutic target. While overexpression using viral vectors is making its way gradually into clinics, CRISPR-mediated targeting of enhancers provides an appealing alternative. Therapeutic enhancer-editing has been demonstrated clinically for therapeutic purposes ^73^ as is the case of the erythrocyte-specific enhancer for the *BCL11A* gene, a transcriptional repressor of fetal hemoglobin^74^. This genome editing leads to the suppression of *BCL11A* in erythrocytes, with a consequent sustained expression of fetal hemoglobin genes and symptomatic relief of sickle cell anemia ^74^. Genome editing of the promoter region of the fetal hemoglobin genes (*HBG1/HBG2*) has also been used to increase the expression of the gamma-globin genes. This led to persistent *HBG1/HBG2* expression and has been proposed as an alternative therapy to the enhancer editing of BCL11A ^75^. However, in this case, given that the promoter region of *LIPTER* is accessible in all cell types, promoter editing may lead to undesirable changes in other cell types. In contrast, the CM specificity of this regulatory enhancer will guarantee increased *LIPTER* expression only in CMs. Ongoing studies will evaluate the use of CRISPR activation (CRISPRa) of the enhancer to increase the abundance of *LIPTER* in hESC-CMs challenged with high concentrations of lipids. We are also identifying key nucleotides and regions within the enhancer which can be edited to increase the abundance of *LIPTER*.

In conclusion, *LIPTER* is a highly abundant CM-enriched lncRNA with an important role in CM development and function. It is conserved in primates and has poor sequence conservation in mice and rats, making *in vivo* studies challenging. However, the recent report showing it has therapeutic relevance is exciting, and our discovery of a CM-specific regulatory enhancer opens avenues to explore gene therapy based on *LIPTER* overexpression.

## Acknowledgements

We thank Azadeh from the Montreal Heart Institute for helping with cloning, and Louis for helping with microscopy.

## Supplementary Figure Legend

**Supplementary Figure 1.**
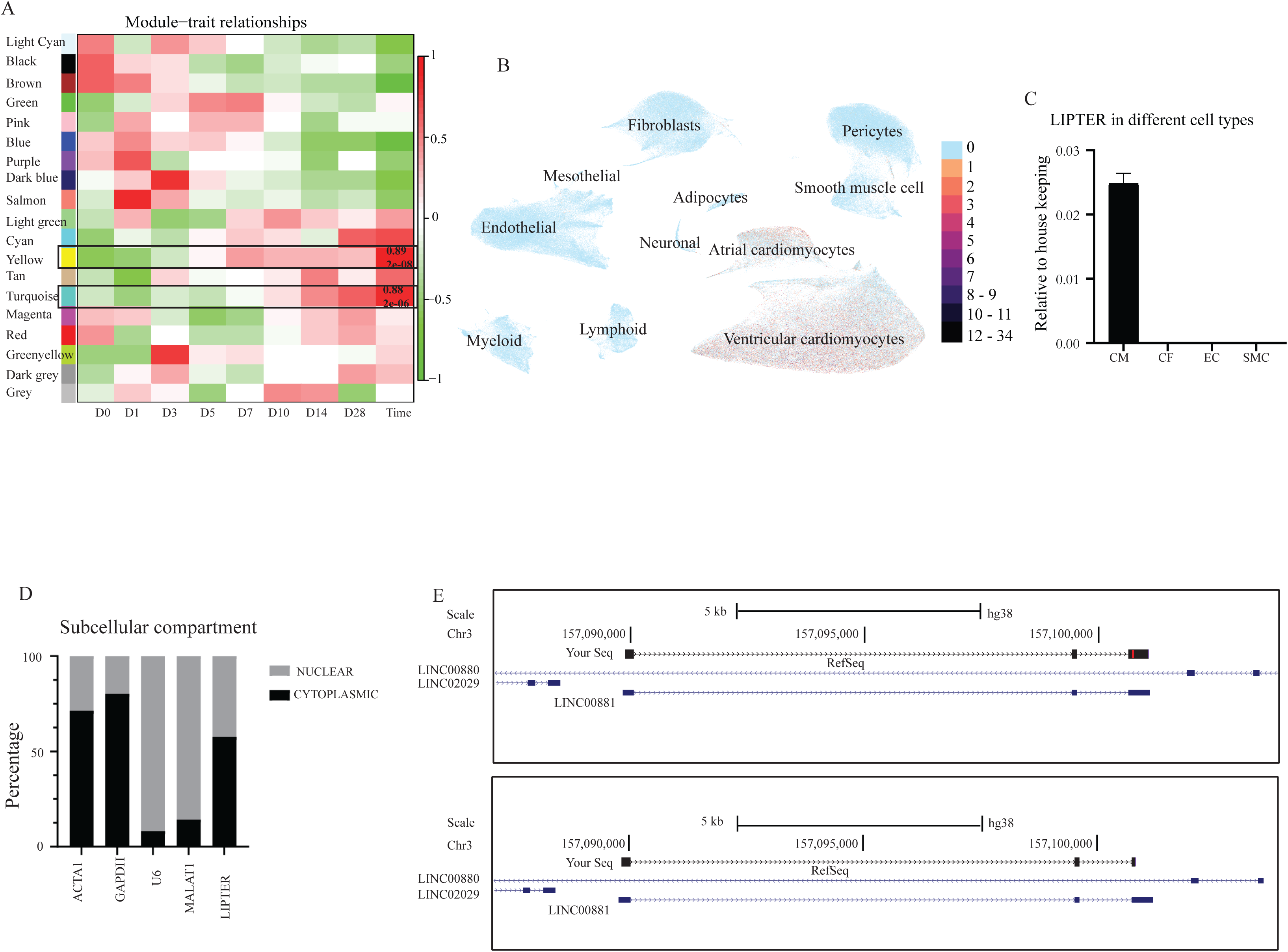
A) Module-Trait relationship from the WCGNA analysis of the RNA-seq data during hESC-CM differentiation. The highlighted modules belong to the fetal genes (yellow) and mature genes (turquoise) B) Umap from a previously published single-cell RNA-seq data^29^ showing the expression pattern of *LIPTER* in the different cell types of the heart. It is only expressed in atrial and ventricular CMs. C) QPCR analysis of *LIPTER* in CM, CFs, ECs and SMCs, showing that *LIPTER* is only expressed in CMs. D) Subcellular fractionation experiment showing higher abundance of *LIPTER* in the cytoplasmic fraction. E) Sanger sequencing result of our rapid amplification of cDNA ends mapped to UCSC genome browser. The most abundant isoform aligns with the RefSeq-identified transcript.

**Supplementary Figure 2.**
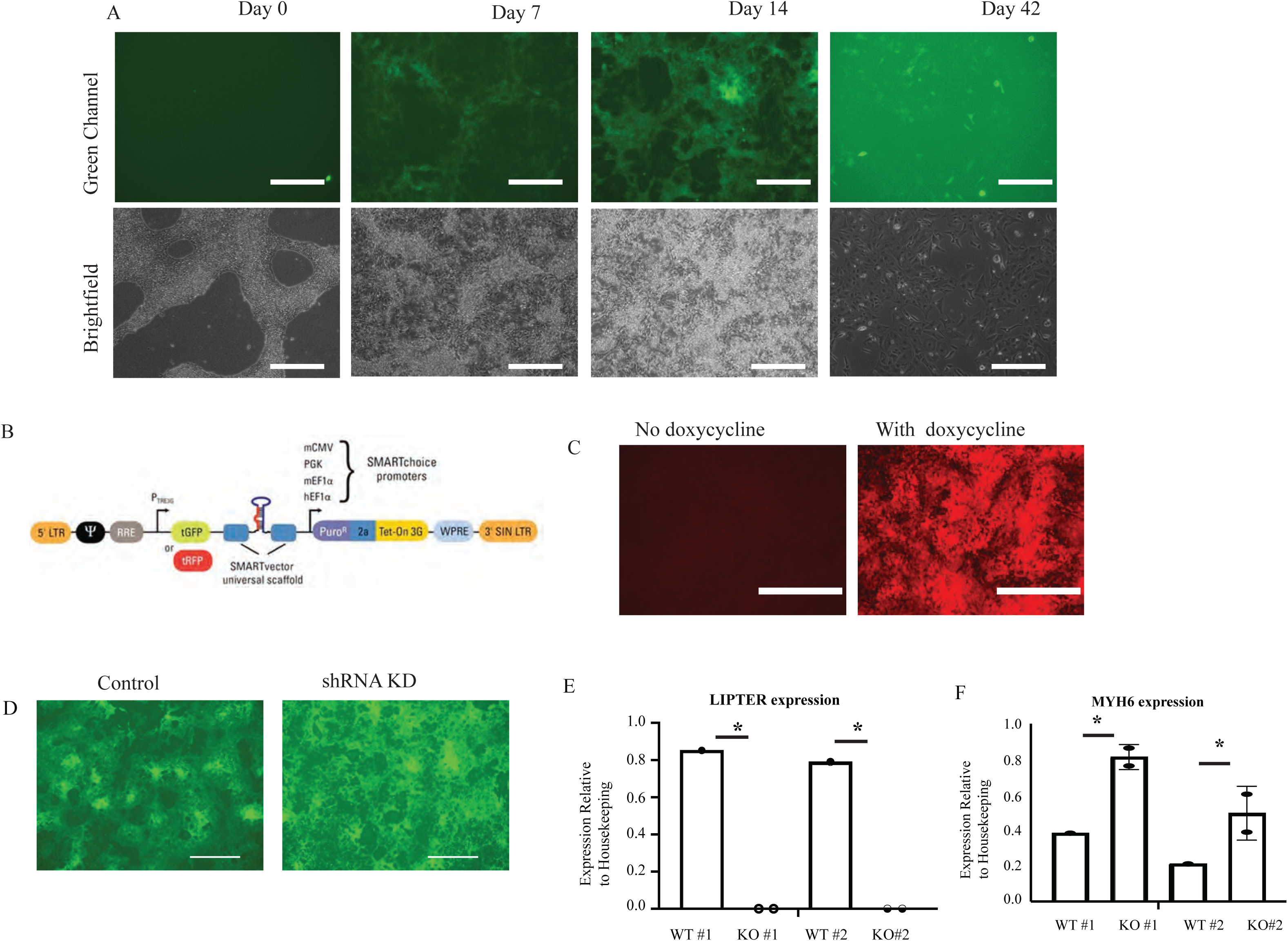
A) Fluorescent images showing the dynamic intensity of CFP indicative of MYH6 during CM differentiation. It is absent in stem cells (day 0), is expressed from day 7 and reduces as CMs mature (day 42). B) A schematic of the shRNA inducible vector. C) Fluorescent images showing RFP intensity indicative of shRNA upon treatment with doxycycline. D) Fluorescent images of day 14 CMs showing similar CFP intensity in control and knockdown cells. E) A qPCR analysis of *LIPTER* expression in control and Knockout CMs. F) A qPCR analysis of *MYH6* in the control and *LIPTER* knockout CMs. Scale bar 50 μm.

**Supplementary Figure 3.**
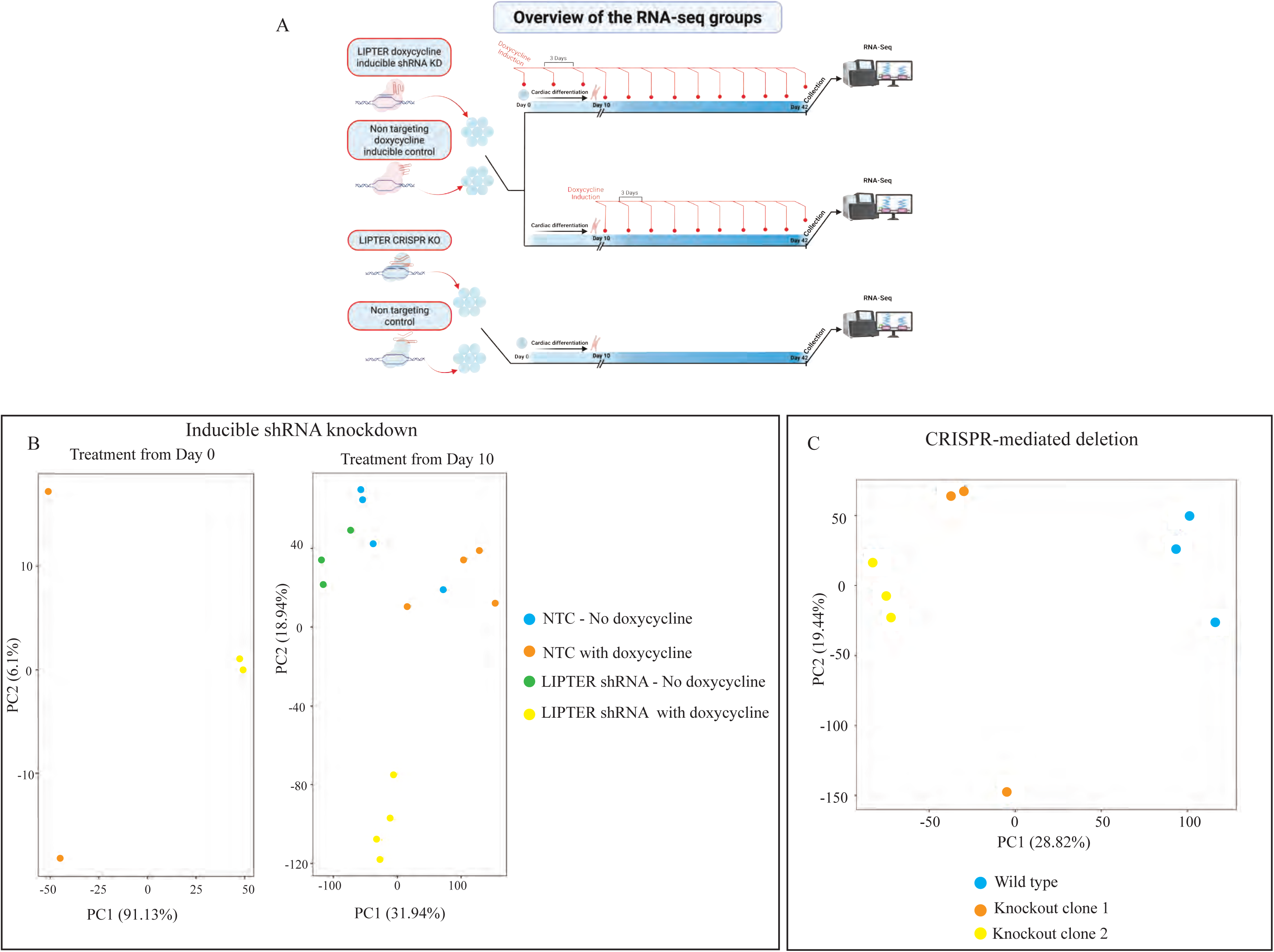
A) Schematic overview of the RNA-seq groups. B) PCA plot showing the clustering of the RNA-seq samples from the inducible shRNA knockdown group. C) PCA plot showing the clustering of the RNA-seq samples from the CRISPR-induced knockout group.

**Supplementary Figure 4.**
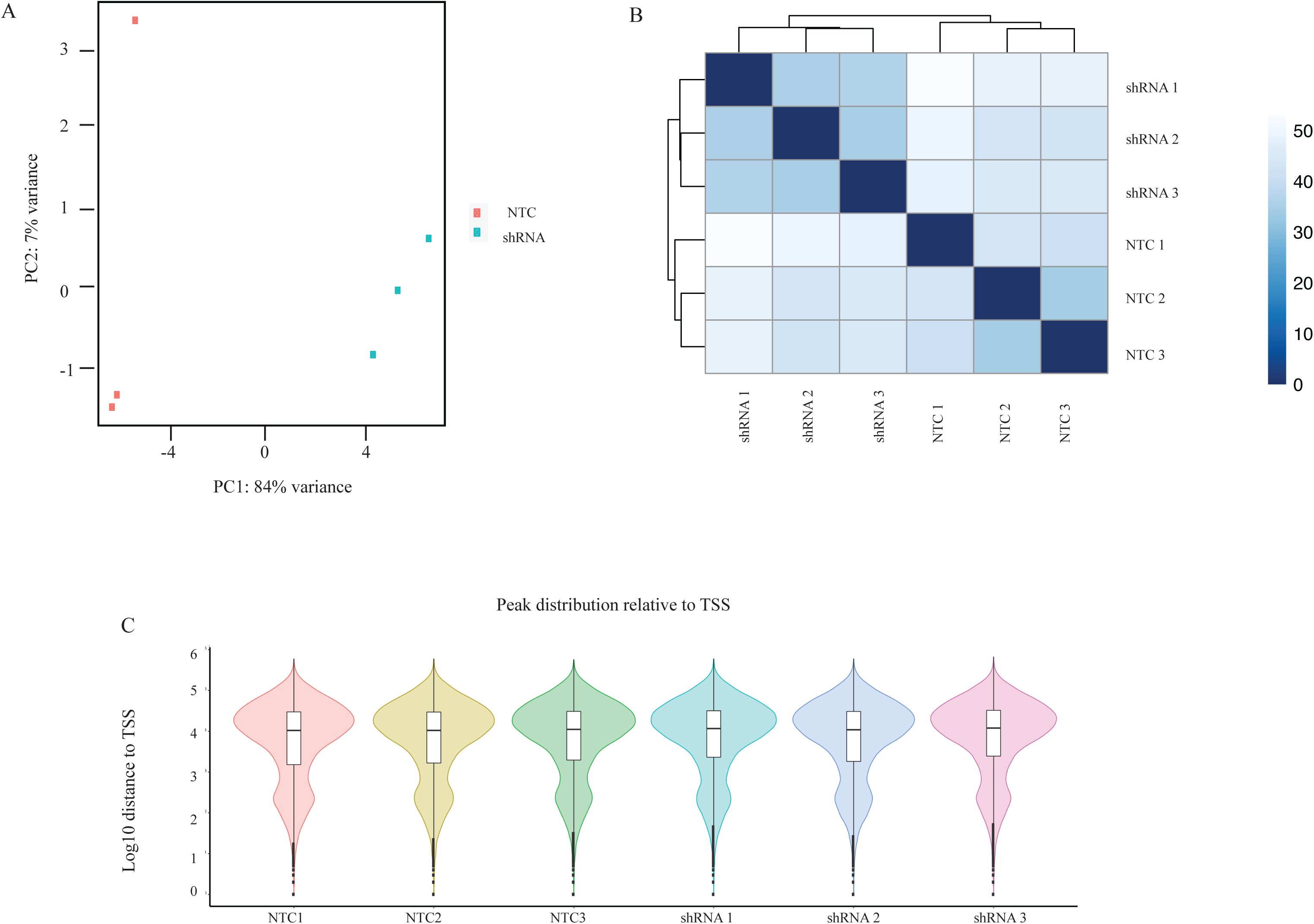
A) PCA plot showing the clustering of the ATAC samples. B) Heat map showing the clustering of the ATAC samples. C) Distribution of the ATAC-seq peaks relative to transcriptional start sites.

**Supplementary Figure 5.**
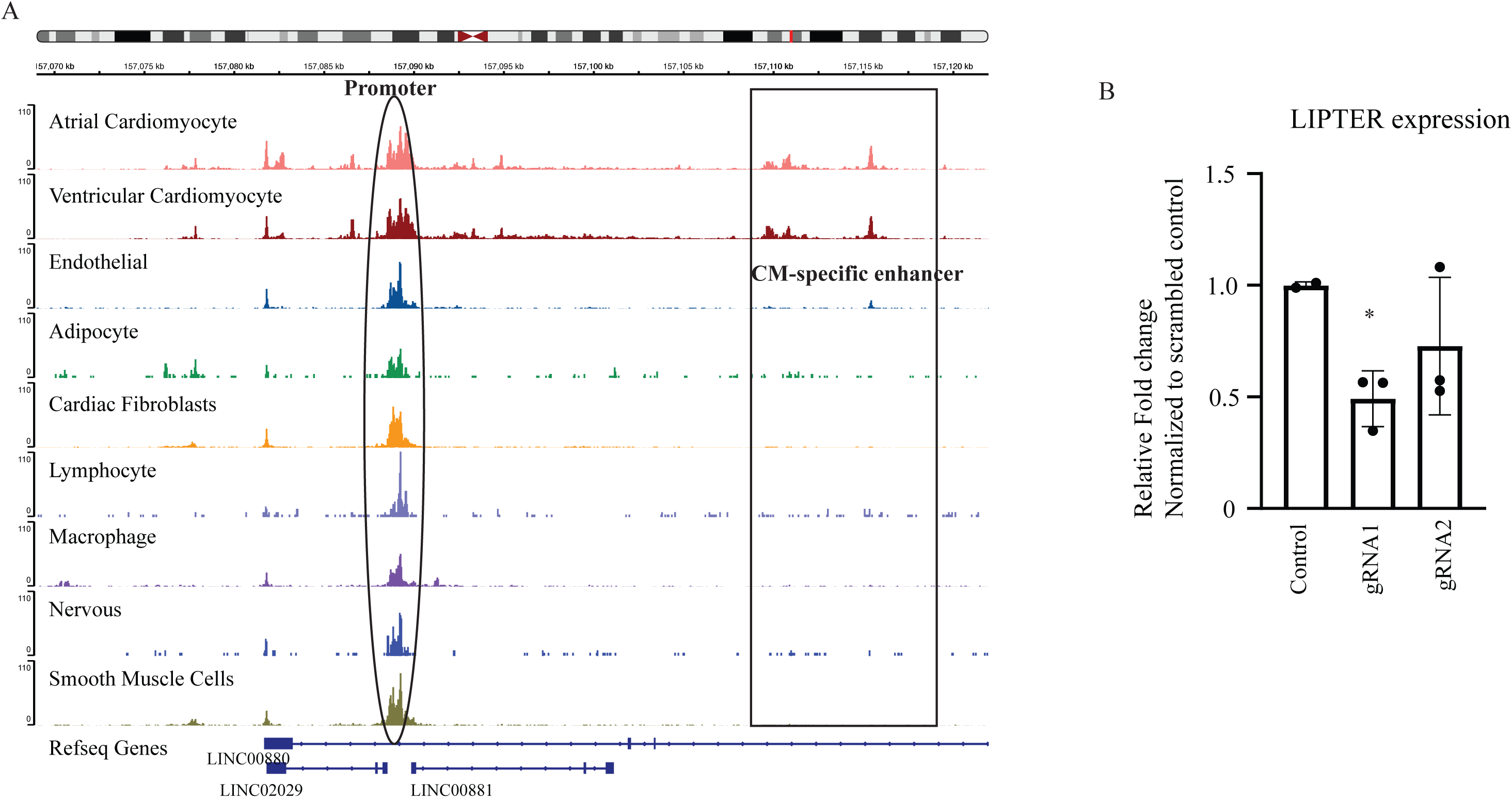
A) IGV plot of previously published single-nuclear ATAC-seq of the heart from CATlas^50^. The black oval highlights the promoter region of *LIPTER* and shows that it is accessible in all cell types of the heart. The grey box shows the cardiomyocyte-specific enhancer linked to *LIPTER*. B) QPCR analysis of the expression of *LIPTER* after enhancer CRISPRi in the hESC-CMs compared to the scrambled control. * P < 0.05

## EXTENDED METHODS

### Research compliance and ethical regulation

All experiments performed with human ESCs using the WA01 (H1) cell lines were under the supervision and guidelines of the National University of Singapore Institutional Review Board (NUS-IRB) committee and the Montreal Heart Institute ethics committee. All mice were maintained and studied under protocols approved by the Institutional Animal Care and Use Committee of the National University of Singapore. Studies were conducted in C57BL/6J mice.

### Human embryonic stem cell differentiation

The cardiomyocyte differentiation was performed using the GiWi protocol adapted from Lian et al^28^. Briefly, the hESC lines were seeded for differentiation on Geltrex-coated plates (A1413202, ThermoFisher Scientific) in mTesR1 and 10 μm ROCK inhibitor (Y-27632) for 24 h. At Day 0, Mesendoderm induction by 6 μM CHIR99021 (#72054, STEMCELL Technologies) in fresh RPMI/B27 without insulin was added for 24h. At Day 3, the media was changed and supplemented with 5 μM IWP2 (I0536, Sigma Aldrich). At Day 5, IWP2 was removed from the media, and RPMI/B27 with insulin (#17504044, Gibco) was added from Day 7 onwards. Cardiomyocyte media was refreshed every 3 days. Metabolic selection using D-lactate was started from Day 14 onwards to remove non-myocyte layers.

### Generation of knockout and knockdown cell lines

The H1 MYH6-mCerulean3 reporter line engineered by knock-in of the mCerulean3 transgene at the 3’ end of the MYH6 endogenous locus was used for all the experiments^31^. The dCas9-KRAB derivatives were generated using lentivirus to stably integrate the dCas9-KRAB complex into the hESC cells^76^.

#### Knockout lines

We performed CRISPR-mediated knockout of the *LIPTER* by excising the promoter and first exon from the genomic locus, using dual single guide RNAs (gRNA). Two independent monoclonal *LIPTER* KO lines were generated from the H1-hESC-MYH6-mCerulean3 reporter line using plasmid pMIA3 (Addgene plasmid No. 109399). Cloned single guide ribonucleic acids (gRNAs) were designed by identifying NGG PAM (protospacer-adjacent motif) sites targeting the promoter and exon 1 of the *LIPTER* gene locus. Briefly, the H1-hESC-MYH6-mCerulean3 reporter line was dissociated using Accutase (#07922, STEMCELL Technologies), and ∼2.0 × 10^6^ cells were pelleted before resuspending with 100 μl of P3 Nucleofection Primary Cell Kit solution (V4XP-3024, Lonza) and 10 μg of pMIA3-LIPTER-sgRNA plasmid. Nucleofection was performed using the program setting CM-113 on the 4D-Nucleofector System. Cells were seeded in Geltrex-coated plates supplemented with 2 ml of CloneR™ (#05888, STEMCELL technologies) per well diluted in mTesR1 (1:10 dilution). 2 days after nucleofection, cell colonies were dissociated into single cells with Accutase and FACS-sorted for RFP-positive cells. Single cell colonies were monitored and maintained in mTesR, individual colonies were allowed to expand and isolated as monoclones. Genomic DNA was extracted for genotyping. After genotyping we selected two clones for further experiments

#### Inducible shRNA knockdown lines

The SMARTvector inducible lentiviral particles for the knockdown of *LIPTER* were purchased from Horizon Discovery through Research Instrument Private Limited Singapore (V3SH7966-13EG100498859) shRNA sequence TGCACCAGGACTGAACAGA. Before transduction into hESCs, the cells were pre-treated with 8 μg/ml polybrene, and a high multiplicity of infection was used. Subsequently, puromycin 1 μg/ml (P9620, Sigma Aldrich) was used to select the cells that integrated the construct before single clones were picked.

#### CRISPRi enhancer knockdown lines

The gRNAs targeting the enhancer were coned into the LentiGuidePuro construct. Briefly, LentiGuidePuro was digested with Esp3I (NEB, #R0734L) and annealed sgRNAs were ligated using T4 DNA Ligase (NEB, #M0202L) following manufacturer’s instructions. Following transformation (NEB 5-alpha, NEB, #C2987), plasmid integrity and sgRNA integration were verified by sequencing. Lentivirus for genomic integration of the sgRNAs were produced in HEK293T cells by co-transfecting the sgRNA-carrying LentiGuidePuro vectors and lentiviral packaging plasmids PMD2.G (Adgene, #12259) and psPAX2 (Adgene, #12260). On the third day after transfection, lentiviral particles were harvested using Lenti-X Concentrator (Takara, #631232) according to the manufacturer’s instructions.

Two monoclonal CRISPRi lines targeting the *LIPTER* enhancer region were developed. We first generated a H1 hESC-dCas9KRAB-MYH6-mCerulean3 line stably expressing the dCas9KRAB protein complex. Using the CRISPICK tool, two sgRNAs targeting the *LIPTER* enhancer region were designed and cloned into the LentiGuidePuro vector (Adgene, #52963). The day before transduction, H1 hESC-dCas9KRAB-MYH6-mCerulean3 cells were seeded onto Matrigel-coated 12-well plates at a density of 150 000 cells/cm^2^ in mTeSR (Stemcell Technologies, #85851) supplemented with 10µM Y-27632 (Stemcell Technologies, #72308). Cells were transduced with lentiviral particles using polybrene at a final concentration of 7.8µg/µL for 24h. Cells were allowed to recover for 72h before puromycin (Gibco #A11138-03) selection at 1µg/mL.

### Patch Clamp

#### Perforated patch-clamp recording of spontaneous action potential

Action potentials from the hESC-derived cardiomyocytes were recorded with external solution containing (in mM)10 Glucose, 125 NaCl, 25 NaHCO_3_, 1.25 NaH_2_PO_4_.2H_2_0, 2.5 KCl, 1.8 CaCl_2_, 1 MgCl_2_, pH 7.4 (300–310 mOsm) and internal solution containing (in mM) 130 K-gluconate, 10 KCl, 5 EGTA, 10 HEPES, 0.5 Na3GTP, 4 MgATP, Na-phosphocreatine, adjusted with KOH to pH 7.4 (290 mOsm). The internal solution was supplemented with 12.5 mM of escin to perform perforated patch clamp.

#### Whole-cell patch-clamp recording of voltage-gated calcium current

To record Ca_V_1.2 current, the internal solution (patch-pipette solution) contained the following (in mM): 138 Cs-MeSO_3_, 5 CsCl, 5.0 EGTA, 10 HEPES, 1 MgCl_2_, 2 mg/ml Mg-ATP, pH 7.3 (adjusted with CsOH), 290 mOsm with glucose. The external solution contained the following (in mM): 10 HEPES, 140 tetraethylammonium methane sulfonate, 1.8 CaCl2 (pH adjusted to 7.4 with CsOH and osmolality to 290-310 with glucose). Pipettes of resistance 1.5-2 MΩ were used. Whole cell currents were obtained under voltage clamp with an Axopatch200B or Multiclamp 200B amplifier (Molecular Devices), low-pass filtered at 1 kHz, and the series resistance was typically < 5 MΩ after > 70% compensation. The P/4 protocol was used to subtract online the leak and capacitive transients.

### Immunocytochemistry

Briefly, CMs are fixed with 3.7% formaldehyde and permeabilized with 0.2% Triton X-100/PBS. Blocking is performed with bovine serum albumin (BSA)/PBS before incubation with mouse anti-cardiac actinin primary antibody (Sigma-Aldrich Cat# A7811) in 10% normal goat serum at 4°C overnight. The next day, the primary antibody is washed off with PBS and incubated with secondary antibody Goat anti-Mouse, Alexa Fluor 594 (Thermo Fisher, cat# A-11005) in 10% normal goat serum for 2 hours, and subsequently with DAPI.

### Sarcomere organization analysis using the SotaTool

The SotaTool software was run following published instructions ^45^. All images used were captured using the same confocal microscope and processed equally. The parameters for the SotaTool were as follows: Resolution + 2 pixels/μm, Background subtraction = True, Segmentation = True (5 x 5). Other parameters were kept at their default settings. Three different images for each group were used for the analysis. Data is presented as a boxplot with the interquartile range. A two-tailed unpaired Student’s t-test was performed for comparison between the two groups. All tests were performed using GraphPad Prism 8, and P < 0.05 was considered significant.

### *In vivo* reporter assay

#### Subcloning of *LIPTER* enhancer fragments for AAV viral vector production

The minimal promoter (minP) from the pGL4.23 (Promega) plasmid was amplified using primers containing 15-30 bp overlaps with the target plasmid and gel-purified. In particular, the forward primer contained an additional *NotI* restriction enzyme cut site. The minP fragment was subcloned into the pENN.cTNT.PI.EGFP.LacZi (Penn Vector Core) plasmid backbone. The plasmid was digested with XbaI (NEB #R0145S) in 1x CutSmart™ Buffer for 1.5 hours at 37 °C before gel purification. Following which, digested vector and insert fragments were added in a 1:3 vector: insert molar ratio to the NEBuilder HIFI DNA Assembly Master Mix (NEB #E2621S) and incubated at 50 °C for 15-60 minutes. 5 uL of the ligation mixture was then used for subsequent transformation into RapidTrans™ Chemically Competent Cells (ActiveMotif #11096), forming plasmid minP-Laczi-GFP. The *LIPTER* distal enhancer (hg38 coordinates chr3:157,108,500-157,110,924) was subcloned into minP-Laczi-GFP. Briefly, the region of interest was amplified using specific primers containing 15-30 bp overlaps with the target purchased from IDT. Following this, minP-Laczi-GFP was digested with *NotI* (NEB #R3189S) and *HindIII* (NEB #R3104S) and gel purified. Similarly, the digested vector and insert fragments were added in a 1:3 vector: insert molar ratio to the NEBuilder HIFI DNA Assembly Master Mix (NEB #E2621S) and incubated at 50 °C for 15-60 minutes. 5 uL of the ligation mixture was then used for transformation to RapidTrans™ Chemically Competent Cells (ActiveMotif #11096). Cloned constructs are summarised in Table 1.

**Table 1:**

| Name | Description |  |
| --- | --- | --- |
| Full enhancer | Full enhancer | chr3:157,108,199-157,110,924 |
| Fragment F1 | 735bp region | chr3:157,108,199-157,108,934 |
| Fragment F2 | 735bp region | chr3:157,1,089,02-157,109,637 |
| Fragment F3 | 670bp region | chr3:157,109,605-157,11,0275 |
| Fragment F4 | 735bp region | chr3:157,110,243-157,111,924 |

#### Adeno-associated virus production and purification

Briefly, recombinant adeno-associated virus (AAV) serotype 9 was produced by a transient triple transfection in HEK293T cells. AAV viral vectors containing the candidate regulatory element upstream of an EGFP or luciferase reporter were co-transfected with helper plasmid pAdΔF6 and plasmid pAAV2/9 (Penn Vector Core). Three days after transfection, cells were harvested, and the virus was purified from cell pellets using Optiprep density gradient medium (Sigma #D1556). Viruses were buffer exchanged to 1xPBS without calcium and magnesium with 100 kDa centrifugal filters (Merck Millipore #UFC910024) and stored at −80 °C. Viral titres were later quantified with qPCR.

#### *In vivo* assessment of enhancer activity

Mice were injected with EGFP reporter containing viruses at 10 days old, avoiding any organs. 3 weeks post-injection, mice were anesthetized with 5% isoflurane and the heart and liver excised. The apex of the heart and a small section of the liver were excised for qPCR and stored in Trizol™ Reagent (Invitrogen #15596018), and the remaining tissue was stored in 4% formaldehyde for immunofluorescence.

#### RNA extraction and qPCR

The excised tissues were lysed in Trizol™ utilizing TissueLyser II (Qiagen). Briefly, 5mm stainless steel beads (Qiagen #69989) were added to a 2 ml SafeSeal tube (Sarstedt #72.695.500) containing the tissue-trizol mixture. Tissues were lysed at 30Hz in 2 30s pulses. RNA extraction was carried out following the protocol from Invitrogen. RNA was quantified using a Nanodrop 2000. 1 μg of RNA was treated with 1 μl of DNASE I (Thermo Scientific #EN0521) according to the manufacturer’s protocol. cDNA conversion with the High-Capacity cDNA Reverse Transcription Kit (Applied Biosystems #4368813) was carried out according to the manufacturer’s instructions. The resulting cDNA was diluted with 10 mM Tris-HCl pH 8 and 1uL (∼2.5ng) was used for qPCR. qPCR was carried out using specific primers targeting GFP and 18s (Supplementary Table 16).

### ATAC-seq

The ATAC-seq was performed as per the previously published Omni-ATAC protocol. Briefly, 50,000 cells were pelleted at 500 g for 5 minutes and subsequently resuspended in 100 µl ATAC-resuspension buffer containing 0.5% NP40, 0.5 % Tween-20 and 0.01% Digitonin. The cells were incubated on ice for 10 minutes, and then 1 ml of cold ATAC-RSB containing 0.1% Tween 20 was used to wash the cells. The nuclei were pelleted at 500 g for 5 minutes, the supernatant was carefully removed, and the nuclei were resuspended in 50 µl transposition mixture containing 25 µl of 2x TD buffer, 16.5 µl of PBS, 5 µl nuclease free water, 0.5 µl of 1% digitonin, 0.5 µl of 10% Tween-20 and 2.5 µl of transposase (Illumina Tagment DNA enzyme 1, Catalogue number 20034198). Transposition was performed by incubating the mixture at 37°C for 30 minutes in a thermoshaker, after 30 minutes, the DNA was extracted using the NEB Monarch® PCR & DNA cleanup kit (Catalogue number T1030L). Libraries were generated by performing PCR using the Illumina/Nextera primers after which Ampure XP beads were used for library cleanup, and the DNA libraries were subsequently sequenced on the Illumina NextSeq platform with 75bp Paired-End sequencing.

### RNA isolation, RT-qPCR and RNA-seq

Cells were lysed directly with Trizol reagent (K182001, Themofisher), and total RNA was extracted using Direct-zol™ RNA Miniprep Kit (R2060, Zymo), following the manufacturer’s protocol. Briefly, 500ng of total RNA was converted using the High-capacity cDNA kit (applied biosystems, #4368813) to cDNA and RT-qPCR was performed on 2.5ng of cDNA input. Quantitative PCR for relative transcript expression was performed on an ABI QuantStudio™ 5 Real-Time PCR System. The average Ct values were calculated using the ΔΔCt method to assess relative gene expression changes. Expression levels of targeted genes were normalized against two housekeeping genes, *18S* and *PPIA*. The list of primers and sequences used in this study are listed in Supplementary Table 16.

RNA quality and yield were assessed using an Agilent Tapestation for quality control. Only RNA samples with RNA integrity number above 7/10 were used for RNA-seq. Total RNA library preparations were prepared using TruSeq Stranded Total RNA Library Prep HMR kit (20020596, Illumina) to deplete ribosomal RNA, followed by appending Illumina TruSeq index adaptors to respective cDNA libraries according to the manufacturer’s instructions. After SPRI bead size selection of cDNA libraries to remove excess adaptors, cDNA libraries were quantified using the Agilent DNA1000 kit (5067-1505, Agilent)

### Cellular fractionation

PARIS kit (AM1921, ThermoFisher) was used for cellular fractionation following the manufacturer’s instructions. Briefly, 300 uL of ice-cold cell Fractionation Buffer with Protease and Phosphate inhibitor cocktail was added to the pellet to disrupt the cell membrane, leaving the nuclear membrane intact on ice for 5-10 minutes. Lysates were centrifuged at 500 x g for 5 minutes at 4°C, and the supernatant was transferred to fresh tubes as cytoplasmic fractions. The remaining pellet is washed with Cell Fractionation buffer and then lysed with 100 μl of ice-cold Cell Disruption Buffer to the nuclear pellet as the nuclear fraction.

### RNA-seq and ATAC-seq Bioinformatic analysis

Paired-end reads were mapped against the human reference genome GrCh38/hg38 using STAR aligner68 with default parameters and based upon gene annotation from GENCODE v38. Mapped gene counts were computed using HTseq-count69 and normalized to Counts per Million (CPM) and Fragments Per kilobase of exon per million mapped fragments (FPKM). The edgeR package was used for differential expression analysis. Differentially expressed genes were called based on average FPKM>1, LogFC>0.5 and FDR<0.05 cut off.

Gene ontology (GO) analysis was performed on differentially expressed genes using the AmiGO 2 web-based tool (http://amigo.geneontology.org/), separately for upregulated and downregulated gene sets identified in each group. GO terms were assessed for biological processes using default parameters, and common enriched themes were identified through comparative analysis across datasets. Cell-type enrichment analysis was performed using the Enrichr platform (https://maayanlab.cloud/Enrichr/), querying the “Human Gene Atlas” database. The top enriched cell types were identified based on adjusted p-values.

Genes were clustered based on co-expression patterns using the WCGNA R package, using the RNA-seq FPKM sample matrix as input, as previously described ^77^. WCGNA performs pairwise relationships among gene transcripts across samples. The network is specified by the adjacency matrix *a _ij_*, which contains the weighted correlations taking on the continuous value from 0 to 1, and is specified by the formula *log*(*a _ij_*) = β × *log*(*s _ij_*), where sij is defined by the coefficient correlation between genei and genej. Construction of signed gene network and identification of modules were performed using R function, blockwiseModules, with the default parameters, including the following: maxBlockSize = 2500, power = 8, networkType = “signed”, TOMType = “signed”, minModuleSize = 30, reassignThreshold = 0. mergeCutHeight = 0.25. A soft thresholding power (β = 8) was used as the lowest power in which the scale-free topology fit index curve reaches a plateau close to the highest R value possible. Briefly, the function pre-clusters nodes into multiple block clusters, using k-means clustering. Each blocks were subjected to hierarchical clustering, and the modules were defined as branches of the dendrogram. For every gene, its expression profile is correlated to the module eigengene of the given module across the differentiation timepoints. Hence, K_ME_ is defined as eigengene-based connectivity or module membership, which takes the continuous value from 0 to 1. The closer the K_ME_ value to 1, the higher the correlation of the gene to the module eigengene.Genes that were highly correlated and co-expressed to the module eigengene (K_ME_>0.75) were used for downstream gene set enrichment analysis, while genes with K_ME_<0.3 were filtered away. All figures were generated using R-3.5.1. The mature gene module was selected based on known markers of CM maturation, namely *MYH7* and *MYL2*, while the early fetal gene module was selected based on known markers such as *MYH6* and *NPPA*.

For ATAC-seq, Paired-end sequencing reads were processed and aligned using the DNA-mapping tool to the hg38 human genome as per default parameters. Cutadapt was used for the trimming of adaptors, and Bowtie2 was used as the aligner. MACS2 was used for peak calling at qval: 0.001 and fdr: 0.05. Genomic Regions Enrichment of Annotations Tool (GREAT) analysis was performed on regions with differential chromatin accessibility. Enrichment for biological process GO terms was calculated using the GREAT web server (http://great.stanford.edu/), and top-enriched terms were selected based on FDR-adjusted significance values. Differential accessibility analysis was performed by quantifying read counts over peak regions and comparing KD versus control samples using DESeq2. Peaks with an FDR < 0.05 were considered differentially accessible.

Motif analysis was done using differential peaks obtained from DESeq2. Differential peaks were defined as peaks with FDR < 0.05 and |log2FC| > 1. Background peaks were defined as (FDR ≥ 0.05 or |log2FC| ≤ 1). Human TF motifs were retrieved from JASPAR2020, motif matching was performed on both differential and background peaks, and motif presence and absence matrices were generated. Fisher’s exact test was conducted for each motif, and the occurrence of the motifs was compared in differential and background peaks. Odd ratios and p-values were calculated, and Benjamini-Hochberg correction was appliedEnhancer–promoter interactions were assessed using the Activity-by-Contact (ABC) model as previously described. Enhancer predictions specific to cardiomyocytes were retrieved.

## DATA AVAILABILITY

All sequencing data related to this manuscript will be deposited in a public repository

